# Deep anatomical and ultrastructural classification of neurons in the zebrafish olfactory bulb

**DOI:** 10.64898/2026.07.13.738197

**Authors:** Nila R. Mönig, Michał Januszewski, Stephan Gerhard, Bo Hu, Nesibe Z. Temiz, Ruth E. Montaño Crespo, Tafheem Masudi, Adrian A. Wanner, Christel Genoud, Rainer W. Friedrich

## Abstract

Neuronal circuits in the olfactory bulb (OB) perform computations fundamental to pattern classification including a decorrelation and normalization of odor-evoked activity. These computations are mediated by diverse interneurons but a comprehensive picture of interneuron types and their microcircuit organization is lacking. We provide a deep anatomical classification of neuron types and their synaptic connectivity in the OB of adult zebrafish, a well-established model to analyze olfactory computations. We reconstructed 459 neurons in an image volume acquired by serial block face scanning electron microscopy and defined 13 neuron classes based on morphological and ultrastructural features. These comprised two classes of projection neurons and 11 interneuron classes, some of which were further separated into subclasses. Ultrastructural information including spine shape, variations in neurite diameter and synaptic arrangements contributed significantly to the distinction of cell types. As in other species, reciprocal synaptic connections were abundant. Targeted synapse annotation revealed systematic connectivity between projection neurons and interneurons. These included microcircuit motifs combining reciprocal and unidirectional connectivity that provide possible structural substrates for gain control and lateral inhibition. The results provide detailed insights into the structural organization of the OB and an anatomical foundation for physiological and computational studies of information processing in olfaction.

## Introduction

The olfactory bulb (OB) is the first processing center for odor information and a model system to study distributed neuronal computations. Sensory input from the nose is transmitted to the OB through an array of discrete glomeruli, each representing a specific odorant receptor and defining a distinct sensory processing channel. In addition, the OB receives top-down input from multiple higher brain areas. Neuronal circuits within the OB transform odor-specific, distributed glomerular activation patterns into spatial-temporal activity patterns that are broadcast to multiple cortical and subcortical targets. These transformations include a decorrelation of activity patterns evoked by chemically similar odors and a normalization of activity evoked by odors of different intensity (Friedrich and Laurent, 2001; Friedrich et al., 2004; Friedrich and Laurent, 2004; Zhu et al., 2013; Gschwend et al., 2015; Roland et al., 2016; Wanner and Friedrich, 2020), thus resulting in a whitening of odor representations. Transformations of odor representations in the OB are modulated by experience (Chu et al., 2016; Yamada et al., 2017; Kudryavitskaya et al., 2021) and may support the formation of memories and internal models by telencephalic networks (Haberly, 2001; Wilson and Sullivan, 2011; Hu et al., 2024). Moreover, conserved anatomical and functional features of the OB may support the consistency of innate olfactory behaviors and perceptions across individuals (Kobayakawa et al., 2007; Dieris et al., 2017).

The neuronal circuitry of the OB is conserved throughout vertebrates (Andres, 1970). Principal neurons (projection neurons; PNs) receive excitatory input from sensory neurons within one or, at most, a few glomeruli. PNs associated with different glomeruli do not make direct synaptic connections but interact via inhibitory interneurons (INs). Two major classes of INs are juxtaglomerular neurons in the superficial layers, which include periglomerular and short-axon cells, and granule cells in the deep layers (Kermen et al., 2013; Nagayama et al., 2014). Each of these IN classes comprises multiple neuron types and is continuously remodeled by lifelong neurogenesis. Synaptic interactions between PNs and INs are often reciprocal, i.e., an excitatory PN-to-IN synapse is often paired with an adjacent inhibitory synapse in opposite direction (Rall et al., 1966).

Despite the conservation of its core circuitry, the OB also exhibits differences between vertebrate classes. In aquatic vertebrates, boundaries of glomeruli are sometimes not clearly delineated by glia (Oka et al., 1982; Satou, 1990; Byrd and Brunjes, 1995a). All vertebrates contain mitral cells (MCs) as a major PN type but additional PNs differ between vertebrate classes. The teleost OB contains a unique PN type, the ruffed cell (RC), that is characterized by dense protrusions at the axon initial segment (the “ruff”) (Kosaka and Hama, 1979; Kosaka, 1980; Kosaka and Hama, 1981; Fuller and Byrd, 2005). The OB of tetrapods contains PNs referred to as tufted cells that are distinguished from MCs by their soma location, projections and physiological properties (Nagayama et al., 2014). In teleosts, MCs are morphologically diverse but it remains unclear whether subtypes of teleost MCs correspond to tufted cells. In addition, the mammalian OB contains an excitatory IN type, the external tufted cell, that has not been described in other vertebrates (Hayar et al., 2004a).

Zebrafish are an important model in systems neuroscience because their small brain enables exhaustive optical measurements of population activity, even at adult stages (Yaksi et al., 2007; Zhu et al., 2012; Huang et al., 2020). Moreover, the small brain size greatly facilitates high-resolution analyses of neuronal morphology and synaptic connectivity (Wanner and Friedrich, 2020; Friedrich and Wanner, 2021). Measurements of odor-evoked activity and behavior in zebrafish provided insights into olfactory information processing in the OB and higher brain areas (Friedrich and Korsching, 1997, 1998; Friedrich and Laurent, 2001; Friedrich et al., 2004; Friedrich and Laurent, 2004; Niessing and Friedrich, 2010; Namekawa et al., 2018; Frank et al., 2019). Moreover, in larval zebrafish, the reconstruction of the full wiring diagram by volume electron microscopy (EM) revealed specific higher-order connectivity that is critical for the whitening of odor representations (Wanner et al., 2016; Wanner and Friedrich, 2020; Friedrich and Wanner, 2021).

The comprehensive reconstruction of neurons in the OB of larval zebrafish revealed a core circuitry consisting of MCs, different types of juxtaglomerular INs, and a few rare cell types (Wanner et al., 2016). However, the largest neuronal class in the adult OB – the granule cells – emerge primarily at later developmental stages. Most functional studies of odor processing were performed in adult zebrafish (Friedrich and Korsching, 1997, 1998; Friedrich and Laurent, 2001; Edwards and Michel, 2002b; Friedrich et al., 2004; Friedrich and Laurent, 2004; Yaksi et al., 2007; Niessing and Friedrich, 2010; Zhu et al., 2013; Kermen et al., 2020) but a comprehensive assessment of neuron types and their connectivity is lacking. More detailed anatomical and structural information is therefore needed to understand mechanisms of olfactory information processing in the adult OB.

We reconstructed hundreds of neurons in a large EM image volume of the adult zebrafish OB and classified neuron types based on morphology and ultrastructure. Detailed anatomical analyses revealed a precise and discrete organization of glomerular structures even in areas of the glomerular layer where sensory neuropil appears diffuse. Morphological analyses identified three subclasses of MCs and eleven distinct IN classes with specific connectivity to PNs. These results provide a comprehensive classification of neuron types in the adult zebrafish OB that delineates synaptic microcircuits and provides a structural basis for mechanistic analyses of neuronal computations.

## Results

### Dataset

We reconstructed neurons in a volume of EM images from the ventro-lateral OB of an adult zebrafish (288 x 173 x 98 μm^3^; voxel size: 9 x 9 x 25 nm^3^; ∼18% of the total OB volume, ∼4500 somata) (Wanner et al., 2016) that was acquired by serial block face scanning EM (Denk and Horstmann, 2004; Titze and Genoud, 2016). The long axis of the volume (288 μm) was oriented radially, spanning all layers, while the short axis (98 μm) extended approximately in anterior-posterior direction (Fig. 1A). The volume targeted a ventro-lateral region referred to as the “lateral chain” (Baier and Korsching, 1994) that is responsive to amino acid odorants (Friedrich and Korsching, 1997, 1998; Yaksi et al., 2007; Yaksi et al., 2009). The volume was located anterior to the characteristic ventro-posterior glomerulus, which is identifiable across individuals (Baier and Korsching, 1994; Braubach et al., 2012a). It overlapped with the region labelled “lG” by Braubach et al. (Braubach et al., 2012b) and possibly contained part of the “lateral plexus” (Baier and Korsching, 1994; Braubach and Croll, 2021).

**Figure 1.**
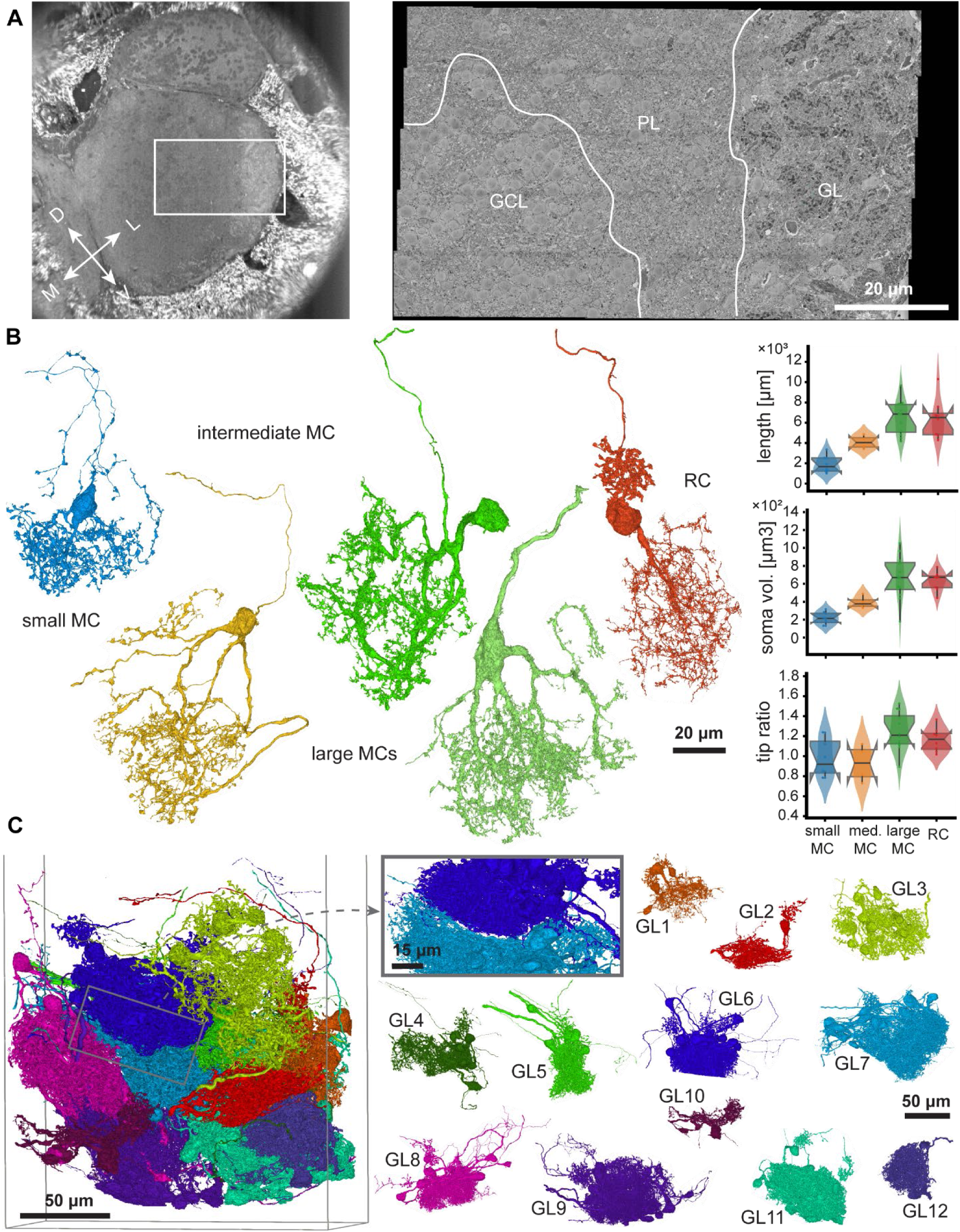
Overview and glomeruli. A. Left: Low-resolution EM image showing a cross-section through the adult zebrafish OB. Box approximates the area where the high-resolution EM volume was acquired. Right: Single high-resolution section in the center of the volume (composite of 40 tiles). Boundaries of the GCL, PL, and GL are outlined. B. Left: Examples of PNs. Right: Quantification of skeletal length, soma size, and the ratio of terminal branches to total dendritic length. Box plots show the distribution of morphological features, where each data point represents an individual neuron within the respective subclass. C. Overlay of PNs showing organization into 12 discrete glomeruli (colors). Left: Anterior view of all reconstructed PNs. Glomeruli are defined by multiple PNs (same color) with overlapping dendritic arbors. Inset: Example showing a precise boundary between two glomeruli (GL6, dark blue, and GL7). Right: All 12 individual glomeruli.

Following previous anatomical descriptions of the teleost OB (Oka et al., 1982; Satou, 1990; Byrd and Brunjes, 1995b) we delineated three layers: (1) the superficial glomerular layer (GL), an intermediate plexiform layer (PL), and the deep granule cell layer (GCL, Fig. 1A). The defining feature of the GL are bundles of varicose sensory axons, characterized by dark cytoplasmic staining (Pinching and Powell, 1971b), that terminated diffusely in subvolumes of diverse shapes. In addition, the GL contained neuropil of postsynaptic neurons and scattered somata. The PL was dominated by neuropil while the GCL contained many somata, often forming clusters.

### Neuron reconstruction

An initial automated reconstruction of neurons was obtained by an oversegmentation of the EM volume using flood-filling networks followed by automated agglomeration (Januszewski et al., 2018). This “first-order” reconstruction had an expected run length (Januszewski et al., 2018) of 741 μm. More complete reconstructions of 459 neurons from all layers (GL: n = 210 neurons; PL: n = 69; GCL: n = 180) were obtained through manual proofreading which added, on average, 19% of the final volume and 35% of the final skeleton length. The 459 neuron reconstructions provide the basis for quantitative analyses. The reconstructions are available in a Google Cloud Storage Bucket (gs://adultob-public) as well as in [neuromaps](https://neuromaps.app/app#!neuromaps/zebrafish_adult_ob1/main.json5). Interactive 3D visualizations can be viewed [here](https://neuroglancer-demo.appspot.com/#!gs://adultob-public/states/all_fully_reconstructed_neurons.json). We provide an in depth description of neuron class morphology via our digital appendix at https://github.com/moenigin/Moenig-2026-zebrafish-adultob.git.

Some neurons could not be fully reconstructed because they extended beyond the boundaries of the EM volume. Morphological completeness was assessed by assigning a “cutting degree” (CD) that ranges from 0 (CD0) to 3 (CD3) to each neuron. Cutting degrees were determined by visual inspection, including comparisons to other neurons with similar morphology, and defined as follows. CD0: All dendritic processes are contained within the EM volume. CD1: Most of the dendritic arborization is contained in the volume except for a minor fraction of distal branches. CD2: A notable fraction of dendrites is cut at the boundaries of the volume, but the reconstruction is expected to include at least 75 % of the total dendritic arbor. CD3: Reconstructions with more substantial cuts.

As neurites of OB neurons can be both pre- and postsynaptic to other neurites it is often ambiguous to classify them as axonal or dendritic based on classsical criteria. Here we refer to projecting neurites with obvious axonal morphology (thin, varicose) and ultrastructural features (presynaptic structures) as axons while all other neurites are referred to as dendrites. Hence, neurites designated as dendrites received not only synaptic inputs but may also make output synapses. We manually annotated synapses between sets of neurons linked to selected glomeruli to obtain a statistical description of connectivity between cell types (Methods, Table 2).

To define morphological neuron types, we first distinguished between PNs (MCs and RCs) and INs. PNs could be clearly identified by their large soma, a projecting axon that was often myelinated, and a compact, highly branched dendritic arbor that received abundant sensory input (Kosaka and Hama, 1982a). RCs were further distinguished from MCs by their characteristic branched protrusions at the axon initial segment (the “ruff”) (Kosaka and Hama, 1979; Kosaka, 1980; Kosaka and Hama, 1981; Fuller et al., 2006b). INs were morphologically diverse and often lacked an axon, as detailed below.

### Glomerular organization of the ventro-lateral OB

The adult zebrafish OB contains ∼140 morphologically diverse glomeruli that are organized into stereotyped regional groups (Baier and Korsching, 1994; Braubach et al., 2012a). Approximately 20% of these glomeruli can be identified across individuals based on their position, shape and size. However, the organization of the dense neuropil in the lateral OB, where the EM volume was acquired, remains poorly understood. While glomeruli have been described in this region, their boundaries are often difficult to delineate and some neuropil regions, particularly the “lateral plexus”, may lack a glomerular compartmentalization (Baier and Korsching, 1994; Braubach et al., 2012a; Braubach and Croll, 2021). It therefore remains unclear to what extent the lateral OB contains glomerular processing channels.

Light microscopic visualizations of PNs led to the description of two types of MCs in the zebrafish OB: *unidendritic* MCs with a single dendritic arbor, and *multidendritic* MCs with multiple primary dendrites. Multidendritic MCs are enriched in the lateral OB (Fuller et al., 2006a) and have been proposed to innervate multiple glomerular structures in this region (Braubach and Croll, 2021). Hence, individual MCs may sample sensory input from different combinations of glomeruli and, thus, mix information across processing channels. Alternatively, multiple dendritic branches of MCs may innervate subcompartments of the same individual glomerulus, thus maintaining a segregation of defined processing channels. Segregated processing channels may also be maintained if distinct sets of MCs innervate consistent combinations of glomeruli. Previous studies could not distinguish between these scenarios because the resolution of light microscopic approaches was insufficient to precisely analyze the innervation of glomeruli by multiple MCs.

To address this issue, we compared ultrastructural reconstructions of all 65 PNs (54 MCs, 11 RCs). We found distinct groups of PNs whose dendrites overlapped extensively but were separated from dendritic volumes of other groups. To corroborate this finding, we coarsely reconstructed 58 additional PNs (50 MCs, 8 RCs) and 15 orphan dendrites that were recognized as PNs based on their morphology (14 MCs, 1 RCs; Methods). The dendritic arborizations of these 138 PNs formed 12 discrete anatomical structures within the image dataset (Fig. 1C, Table 1, [view](https://neuroglancer-demo.appspot.com/#!gs://adultob-public/states/glomeruli.json)). Boundaries between different dendritic volumes were very sharp with virtually no overlap. We therefore refer to the dendritic volumes of these PN groups as glomeruli. This distinct compartmentalization of the neuropil was not obvious in the raw EM image data.

100 of 104 MCs innervated a single glomerulus and the remaining four MCs extended only minor parts of their neurites into secondary glomeruli. Hence, MCs were strictly uniglomerular despite their multidendritic morphology, indicating that segregated glomerular processing channels are maintained in the diffuse neuropil of the lateral OB. Within glomeruli, MC dendrites were often partially aligned with dendrites of other MCs and did not necessarily visit the entire glomerular volume, consistent with glomerular subcompartments described in the mammalian OB (Kim and Greer, 2000).

### Morphological classification of neurons

To establish a taxonomy of neuron types in the adult zebrafish OB, we defined 13 morphological neuron classes based on comparisons across the 459 reconstructed neurons. Classification was performed by human experts who had access to the high-resolution volumetric reconstructions of all neurons and to the EM volume. Efficient comparisons between large numbers of neurons were enabled by a software interface based on neuroglancer (Maitin-Shepard, 2020) that allowed for fast and interactive visualization of arbitrary combinations of neurons in 3D along with EM images (Methods). PNs were divided into two major categories, MCs and RCs, following previous literature (Kosaka and Hama, 1979, 1982a; Fuller and Byrd, 2005; Fuller et al., 2006a; Braubach and Croll, 2021). INs were divided into three major categories based on their distribution across layers. Coarsely reconstructed neurons were not used to define neuron types but classified post-hoc. 11 INs could not be unambiguously assigned and were excluded from classification. Below, we provide detailed descriptions of neuron classes within each category with a focus on INs. Links to interactive visualizations in neuroglancer are provided for each neuron type.

### Projection Neurons

#### Mitral Cells

Somata of MCs (n=104, including 50 coarsely reconstructed neurons, [view](https://neuroglancer-demo.appspot.com/#!gs://adultob-public/states/MC_subclasses.json)) were large (ø: 8 - 24 µm, *x̅* ± σ: 12.4 ± 2.5 µm) with a single primary cilium. Unlike in mammals but consistent with other teleosts (Satou, 1990), somata were distributed throughout the GL, surrounding the glomeruli. Axons, when identified, arose from the soma (n=46) or from a large proximal dendrite (n=24). MCs typically had multiple primary dendrites that branched asymmetrically and formed a dense and complex arbor. Dendrites varied substantially in diameter. Distally, thin dendritic segments alternated with large varicosities (Fig. 2A) that often harbored inbound and outbound synapses. Distal dendrites often terminated in bulbous structures that received input synapses from sensory axons (Fig. 2A). Two classes of synapses were distinguished: Synapses with large vesicle clouds were found primarily in distal varicosities whereas synapses with small densities and vesicle clouds were located primarily in primary dendrites.

**Figure 2.**
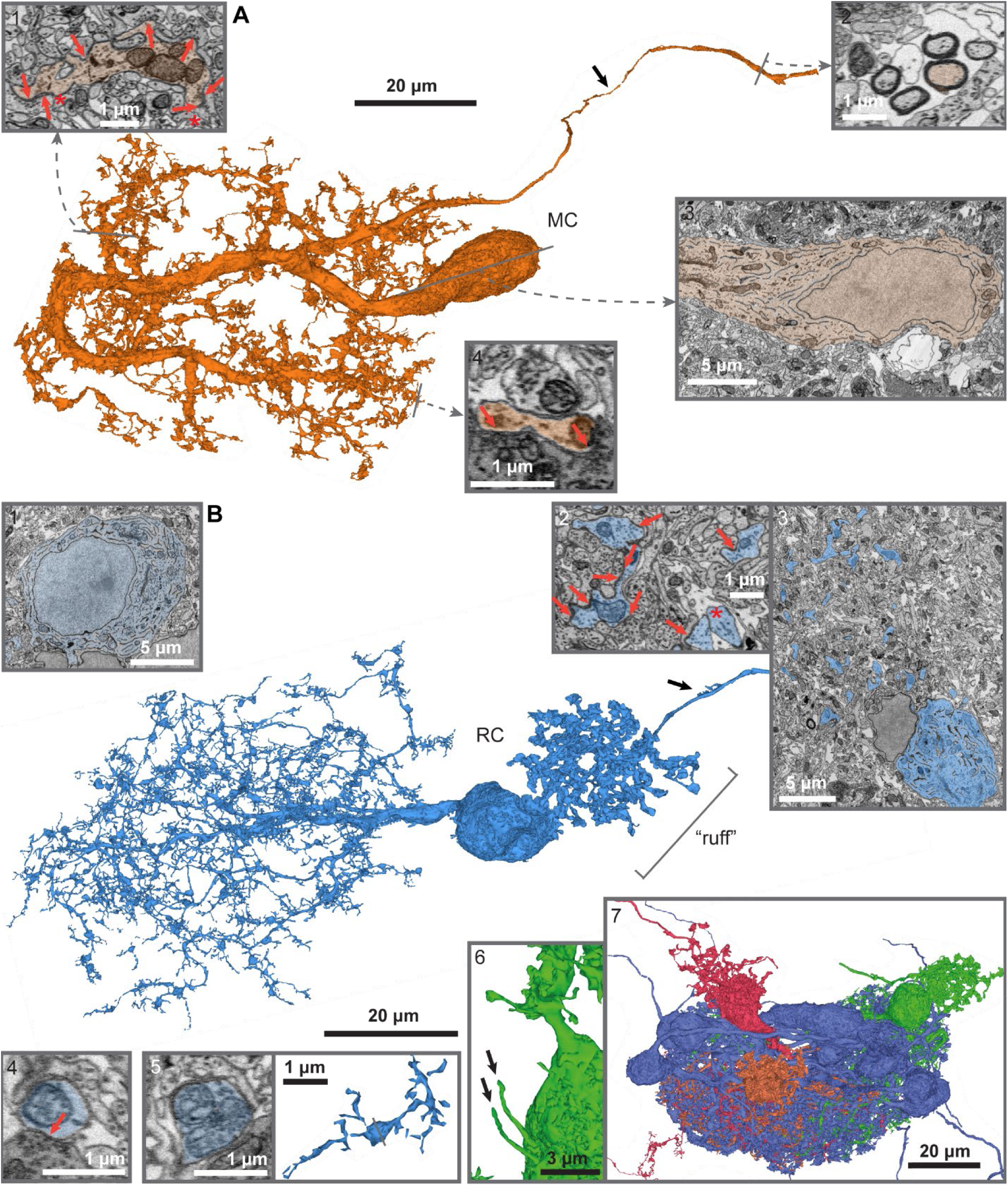
Mitral and ruffed cells. A. Morphology of a large MC. Insets: 1. The MC dendrite exhibits (1) Example of frequent, irregular varicosities containing mitochondria. These enlargements are sites of inbound and outbound synapses with local INs (arrows highlight synapses; asterisks indicate reciprocal synapses). (2) Myelinated MC axons forming small, loose bundles. (3) Irregular surface of a MC soma. The perikaryon contains an abundance of small mitochondria, vacuoles, and endoplasmic reticulum (ER); the Golgi apparatus frequently extends far into the primary dendrite. (4) MC dendrite receiving dense sensory input (arrows) on terminal dendritic branches. B. Morphology of a RC. Insets: (1) RC soma with labyrinth-like ER structures around the nucleus and a multitude of organelles within the endomembrane system. (2,3) Cross sections through the ruff showing RC branches embedded in neuropil that are sites of prominent synaptic input and output, including reciprocal synapses (arrows). The axon (*) emerges from the soma, contains large vesicular structures and lacks obvious outbound synapses. (4) Input synapse from a sensory axon (arrows), which are consistently observed but less frequent than in MCs. (5) Typical thin RC dendrite with a bulbar enlargement containing a few small, round mitochondria with a light appearance. (6) Two primary cilia projecting parallel to the soma surface. Similar structures were observed in approximately half of all RCs. (7) GL6 with three highlighted RCs, showing the location of the somata and ruffs relative to the glomerular neuropil.

MCs were further subdivided into large MCs (n=37, including 18 coarsely reconstructed neurons), intermediate MCs (n=36, including 19 coarsely reconstructed neurons) and small MCs (n=31, including 10 coarsely reconstructed neurons; Fig.1B, Supplementary). Large MCs were characterized by a dense, bushy dendritic arbor and a rough, craggy surface bearing irregular excrescences. Dendrites of intermediate MCs were smoother and less branched. Small MCs had axon collaterals with filamentous protrusions that carried output synapses.

#### Ruffed Cells

RCs (n = 19, including 8 coarsely reconstructed neurons, [view](https://neuroglancer-demo.appspot.com/#!gs://adultob-public/states/RC.json)) had medium-sized, globular somata (ø: 12 - 16 µm, *x̅* ± σ: 14.1 ± 1.4 µm) that were located deep to their parent glomerulus. Most RCs had two long primary cilia (1.8 – 6.5 µm, *x̅* ± σ: 3.9 ± 1.3 µm, Fig. 2B). The characteristic ruff consisted of dense, branched protrusions within the first 5 µm of the axon that extended for 25 - 35 µm into the extra-glomerular neuropil of the deep GL and PL (Fig. 2B, bottom).

Primary dendrites of RCs branched frequently, giving rise to dense, often asymmetric dendritic tufts within the glomerular neuropil. Higher-order branches were thin (∼100 – 200 nm) and twisty with short terminal twigs, contributing a large fraction of the total dendritic path length. While early EM studies in goldfish suggested that RCs receive little or no direct sensory input (Kosaka, 1980), we observed that synapses of sensory axons frequently target the thin, terminal twigs (Fig. 3A, left). The same sensory axons also made synapses onto MCs (Fig. 3A). All glomeruli contained at least one RC.

**Figure 3.**
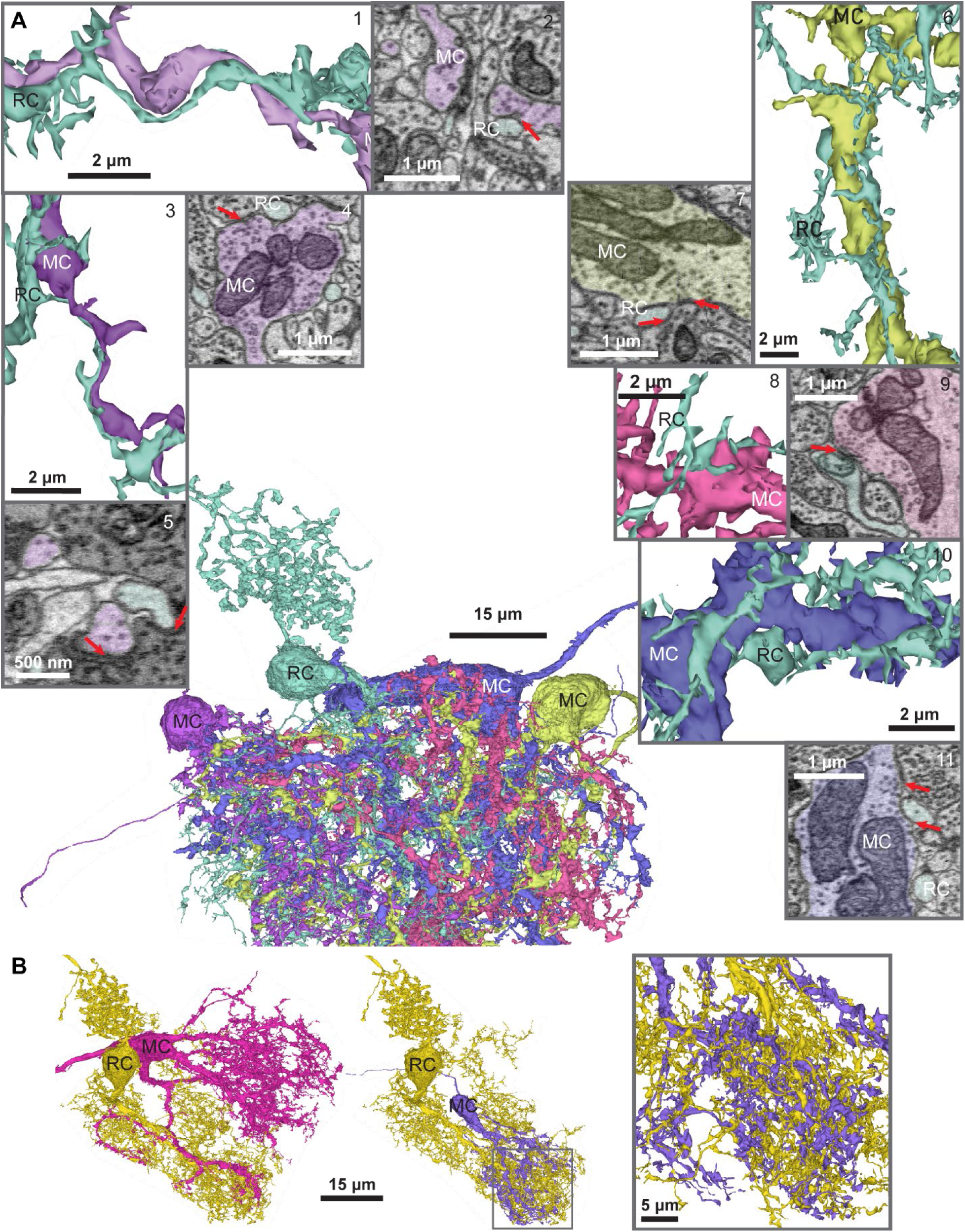
Association between RCs and MCs. A. Center: RC within GL6 and four associated MCs. Arrows highlight synapses. The thin RC dendrites follow and wrap around the MC dendrites (1, 3, 6, 8, 10). Synapses from MCs onto RCs are rare, typically small, and characterized by low anatomical confidence (2, 11), though larger synapses are occasionally observed (9). Terminal branches of the RC often protrude into the space adjacent to synapses between MCs and INs (4, 7, 11), sometimes receiving direct input from the respective IN (7). MCs and RCs receive shared sensory input (5). B. Intertwined dendrites of a RC and two MCs. Inset shows enlargement of boxed area.

Dendrites of RCs and MCs can be intertwined, as observed in ultrastructural reconstructions of small dendritic segments in the goldfish OB (Kosaka and Hama, 1981). Individual RCs formed such dendritic alignments with most MCs within a glomerulus. Although dendrites followed each other for distances up to several micrometers, we detected very few synapses between mitral and RCs (Fig. 3A), often with low confidence. We further observed that physical contacts between a MC and a RC were often located in close proximity to synapses between the MC and an IN (Fig. 3A, right).

### Interneurons

#### Glomerular layer interneurons (GLINs)

##### GLIN1

*Morphology.* GLIN1s (n = 70, [view](https://neuroglancer-demo.appspot.com/#!gs://adultob-public/states/GLIN1.json)) were axon-bearing INs with small somata (ø: 7 - 9 µm, *x̅* ± σ: 7.6 ± 0.5 µm) in the deep GL or superficial PL, often forming small clusters of 3 – 5 neurons near a RC. The thin axons arose from the soma (63/68, unidentified in two GLIN1s) or the proximal dendrite (5/68). Axons emitted collaterals in the GL that passed through one or two glomeruli (*x̅* ± σ: 1.1 ± 0.7 glomeruli) and had prominent varicosities forming outbound and reciprocal synapses with local INs (Fig. 4B).

**Figure 4.**
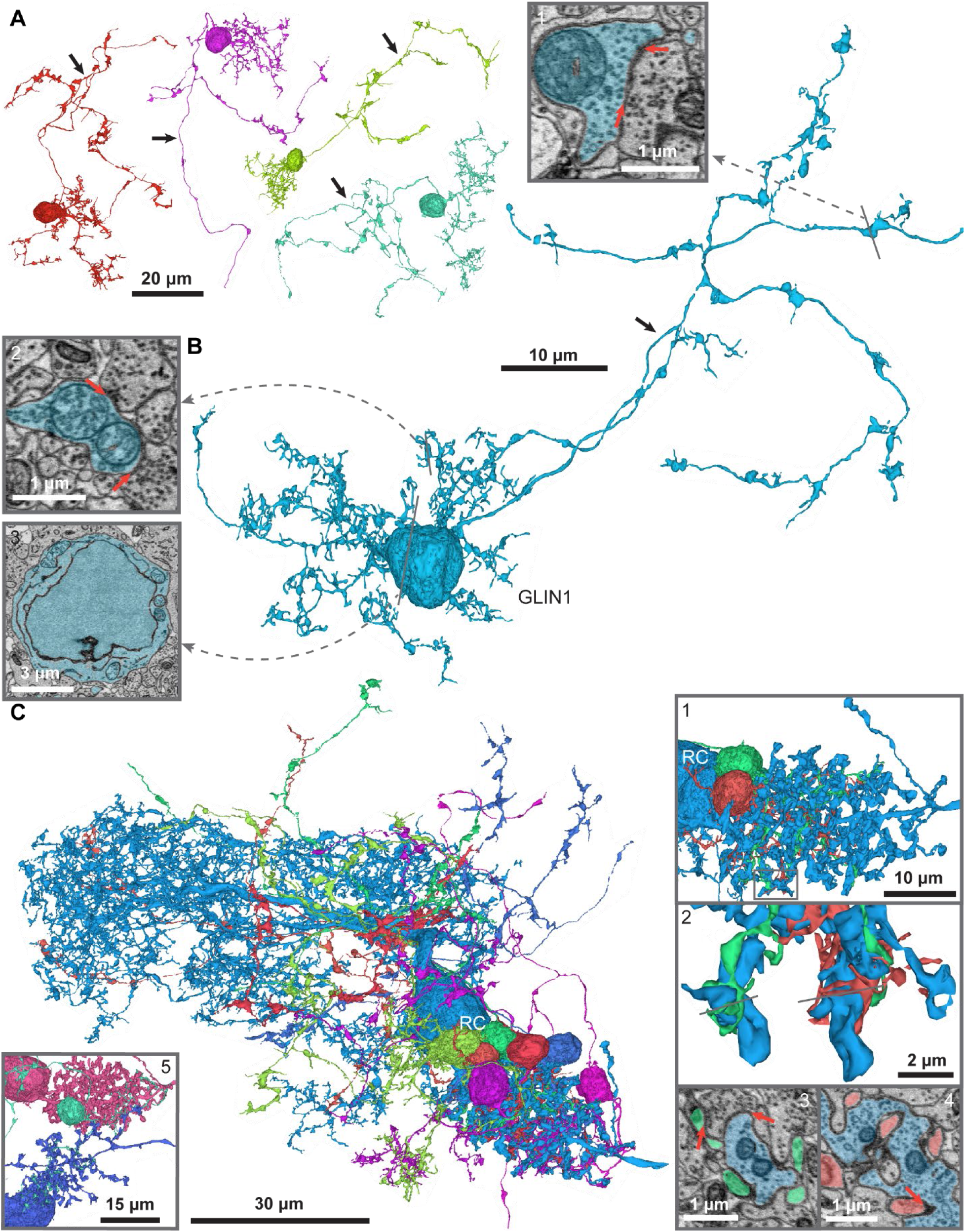
GLIN1. A. Four GLIN1s (arrows: axons). B. Representative example of a GLIN1 with a branched axon (black arrow). Axonal branches contain amorphous varicosities with mitochondria, outbound synapses, and occasional inbound synapses, including reciprocal arrangements (1, arrows). The bushy dendrite, located close to the soma, has frequent bulbar enlargements with mitochondria (2). The soma (3) is small and surrounded by a thin layer of cytosol characterized by small, round mitochondria with a light matrix (see also 2). C. Seven GLIN1s associated with an RC of GL7. GLIN1 dendrites are entirely immersed within the ruff neuropil (1). GLIN1 dendrites tightly wrap around (2) and protrude into (4) RC neurites. Synapses from the ruff onto GLIN1s are often small (arrow in 4). GLIN1s receive input from synaptic partners of the RC (3). Half of the reconstructed GLIN1s interacted with multiple RCs (5).

Up to four short, thin dendrites (ø: 300 - 500 nm) emerged from the soma and formed a bushy tree that was associated with the ruff of a RC (Fig. 4C, Supplementary) and avoided the glomerular volume formed by PNs. No synaptic vesicles and only inbound synapses were observed in the dendrite.

*Interactions with PNs.* The most characteristic feature of GLIN1s was their association with RCs. GLIN1 dendrites wrapped around protrusions of ruffs (Fig. 4C). Despite the large contact area we found only few small but distinct synapses from RCs onto GLIN1s (usually >3 synapses/pair). GLIN1 were primarily associated with a single RC but secondary contacts with a second or third RC (*x̅* ± σ: 1.7 ± 0.7 RCs/GLIN1, Fig. 4B) from the same or different glomeruli were observed in ∼50% of GLIN1s. RCs were associated with up to 14 GLINs. On average, individual GLIN1s were associated with 1.4 ± 0.6 glomeruli (*x̅* ± σ).

Inspections of all 121 GLIN1-RC pairs identified six synapses from the GLIN1 axon onto an RC dendrite or soma (1 – 2 synapses/pair). Five of these connections were reciprocated by input from the RC ruff onto the GLIN1 dendrite. None of the 629 inspected GLIN1-MC pairs were connected.

##### GLIN2

*Morphology.* GLIN2s (n=30, [view](https://neuroglancer-demo.appspot.com/#!gs://adultob-public/states/GLIN2.json)) were distinguished by filiform spine-like appendages. Somata (ø: 8 - 11 µm, *x̅* ± σ: 9.8 ± 0.7 µm) were located in the GL and superficial PL. A thin axon arose from the soma (14/30) or dendrite (11/30) with occasional collaterals and bulbar varicosities that were presynaptic and occasionally postsynaptic to other neurons (Fig. 5B). One or two larger (ø: 0.8 – 3.5 µm, *x̅* ± σ: 1.5 ± 0.7 µm) and up to four smaller primary dendrites (*x̅* ± σ: 380 ± 110 nm) gave rise to many fine higher-order branches, mostly confined to a single glomerulus. Varicosities on small branches projected into the extra-glomerular neuropil (24/30 GLIN2s) and received input from remote axons (Fig. 5B). Higher-order dendrites were covered with hand-like appendages consisting of a very thin and usually long neck that terminated in an enlargement (“palm”) with thin, filopodial protrusions (“fingers”, Fig. 5B). Protrusions usually wrapped around MC neurites while the “palm” received synaptic input.

**Figure 5.**
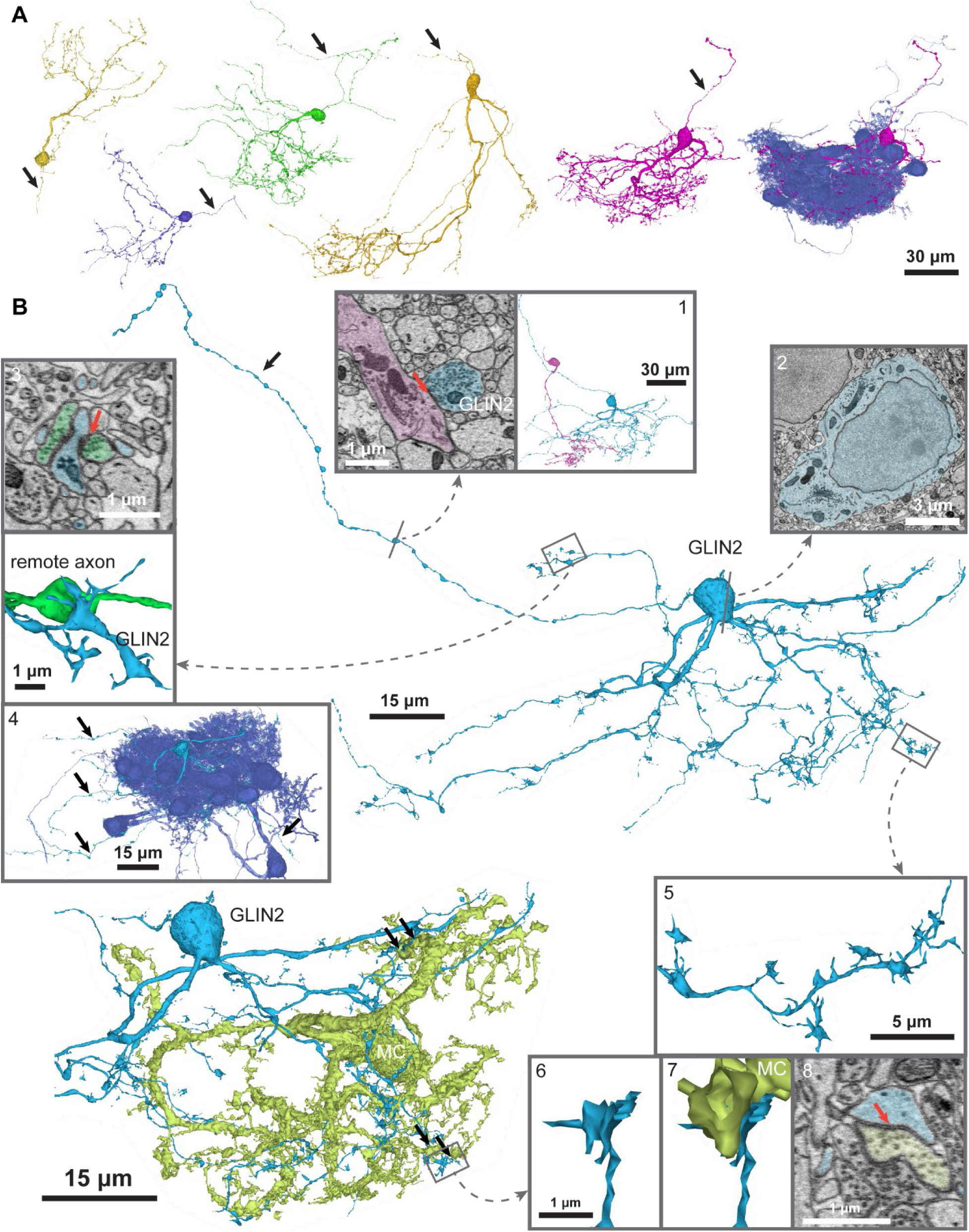
GLIN2. A. Five GLIN2s (arrows: axons) highlighting variable morphology reflecting the shape of the innervated glomerulus (right). B. Example of a GLIN2s with a beaded axon (arrow) extending far into the PL and forming synapses with local INs. Somata are smooth with only occasional small protrusions. (1) Synapse onto a PLIN2 of a neighboring glomerulus (arrow). (2) Nucleus embedded in a prominent layer of cytosol containing relatively little ER and compact, darkly stained mitochondria. (3,4) Minor dendritic segments projecting outside the glomerular neuropil (black arrows in 4; observed in 83% of GLIN2s). These segments form large amorphous structures targeted by axons originating from outside the imaged volume (3). (5) Close-up of the dendrite showing typical spine-like protrusions with a very thin and long neck and a large head with multiple filiform extensions. Bottom left: Association of a GLIN2 with a MC. (6,7,8) Synaptic input from the MC to the GLIN2 via multiple synaptic contacts (arrows) targeting the filiform spine heads.

*Interactions with PNs.* The connectivity between GLIN2s and PNs was examined for five GLIN2s in glomeruli GL6 and GL7. Dendrites of GLIN2s received prominent synaptic input from most or all MCs of their parent glomeruli (often >4 synapses/pair, maximum: 13). No connections between GLIN2s and RC dendrites were found. An additional overlay of 19 GLIN2s with 17 RCs from other glomeruli revealed two synapses from a RC onto the dendrites of two GLIN2s.

##### GLIN3

*Morphology.* GLIN3s (n = 18, [view](https://neuroglancer-demo.appspot.com/#!gs://adultob-public/states/GLIN3.json)) are axonless and characterized by a hairy proximal dendrite and a bushy, beaded distal dendrite (Fig. 6A). Their somata (ø: 8 - 10 µm, *x̅* ± σ: 8.8 ± 0.6 µm) were located in the deep GL or PL. Most GLIN3s (15/18) had a single main dendrite (ø: 0.5 – 1.5 µm, *x̅* ± σ: 1.1 ± 0.4 µm) that projected into a single glomerulus. Five GLIN3s projected a small portion of their dendrite into a second glomerulus or outside the EM volume (mean ± σ: 1.2 ± 0.4 glomeruli). 12 GLIN3s gave rise 1 – 3 additional fine dendrites (ø: 300-500 nm).

**Figure 6.**
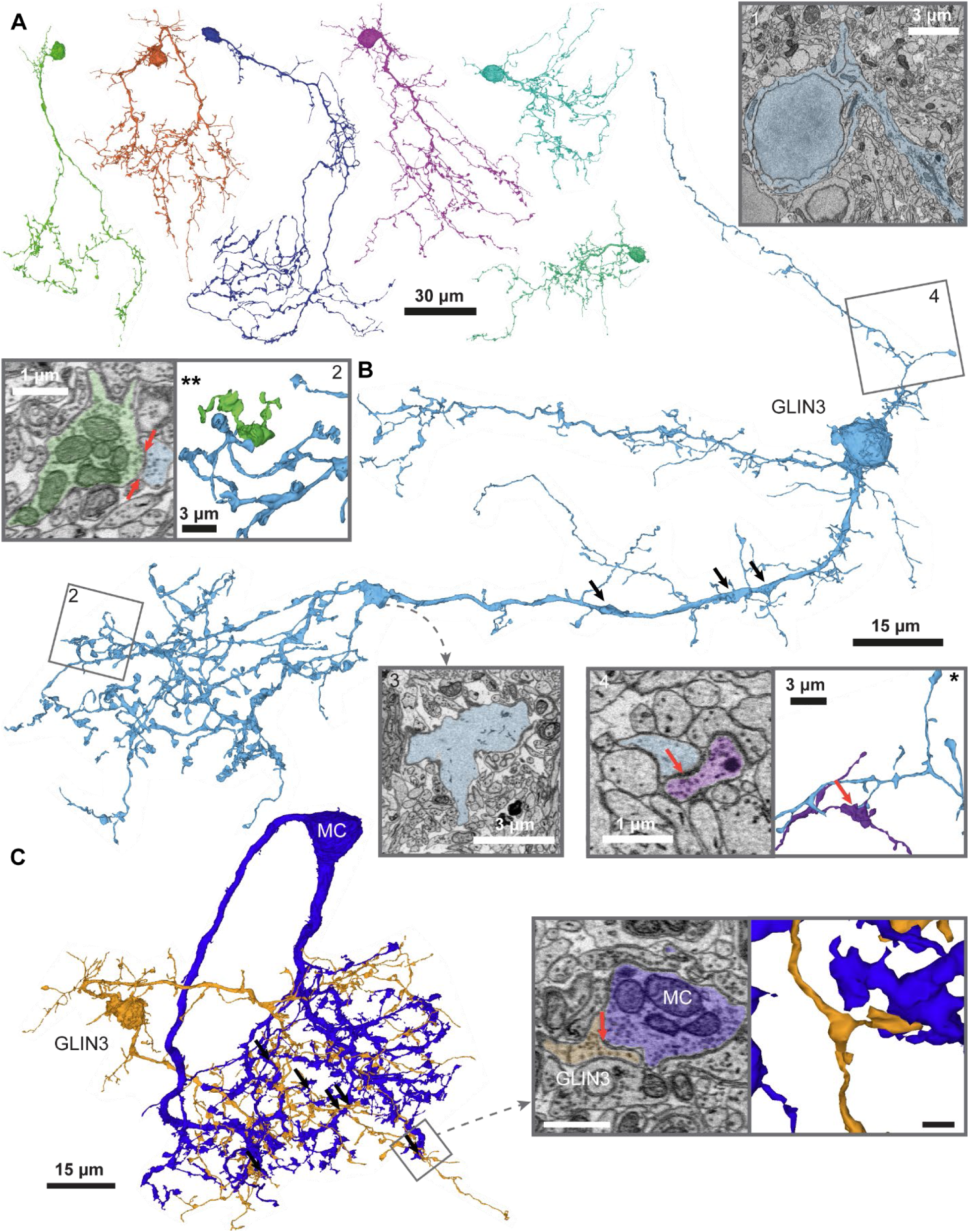
GLIN3. A. Four large and two small GLIN3s. B. Representative GLIN3 with an uneven, craggy surface bearing excrescences and protrusions, as observed in a subset of GLIN3s (e.g., orange and purple GLIN3s in A). The nucleus is surrounded by sparse cytosol. The Golgi apparatus is frequently located within bulges of the primary dendrite at a distance from the soma (1). Proximal dendrites are hairy, covered with thin branches and long spine-like protrusions that are exclusively postsynaptic (4). The primary dendrite exhibits regular enlargements (3; black arrows). (2) Close-up of the distal dendrite showing spine-like varicosities forming reciprocal synapses with MCs (arrows). C. GLIN3 that is reciprocally connected to a MC by multiple synapses in GL6. Inset: Synapse from the GLIN3 onto the MC (arrow).

Proximal dendrites were covered with thin higher-order branches of variable length and long spine-like appendages, resulting in a hairy appearance, and received many synaptic inputs (Fig. 6B) but no output synapses were found. The main dendrite branched and assumed a bushy appearance as it entered the glomerular neuropil. These distal dendritic portions were studded with coin-shaped or varicose enlargements that formed incoming, outgoing and reciprocal synapses with multiple other neurites (Fig. 6B).

*Interactions with PNs.* Synaptic connectivity was assessed between six GLIN3s and PNs in four glomeruli (GL6:3, GL7:1, GL11:1, GL12:1). GLIN3s connected to at least 25 % of the MCs within a glomerulus (ID414: 4/8 MCs (50 %), ID987: 7/12 MCs (58 %), ID1330: 3/12 MCs (25 %), ID1008: 6/7 MCs (86 %), ID1346: 5/12 MCs (42 %), ID2868: 4/12 MCs (33 %)). Most pairs (24/29) were connected through more than three synapses. Most pairs (23/29) were reciprocally connected, usually by reciprocal synapses, but unidirectional GLIN3-to-MC (2/29) and MC-to-GLIN3 connections (4/29) were also found. One GLIN3 formed synapses onto the dendrites of all three RCs of GL6 (3/16 RC-GLIN3 pairs).

##### GLIN4

*Morphology.* GLIN4s (n = 16, [view](https://neuroglancer-demo.appspot.com/#!gs://adultob-public/states/GLIN4.json)) were similar in appearance to GLIN1s with small, typically ovoid somata (ø: 7 - 9 µm, *x̅* ± σ: 8.5 ± 0.6 µm) in the GL or PL. Axons arose mostly from the soma with collaterals traversing one or two glomeruli (*x̅* ± σ: 1.1 ± 0.3 glomeruli, Fig. 7A, B). Axons spread in the GL and PL and formed bundles with GLIN1 axons. GLIN4s had a single thin dendrite (ø: 0.3 - 1 µm) that emerged from the soma and, after tens of microns, gave rise to a sparse, bushy arbor that innervated one (14/16) or two glomeruli (2/16). No outbound synapses were detected in GLIN4 dendrites.

**Figure 7.**
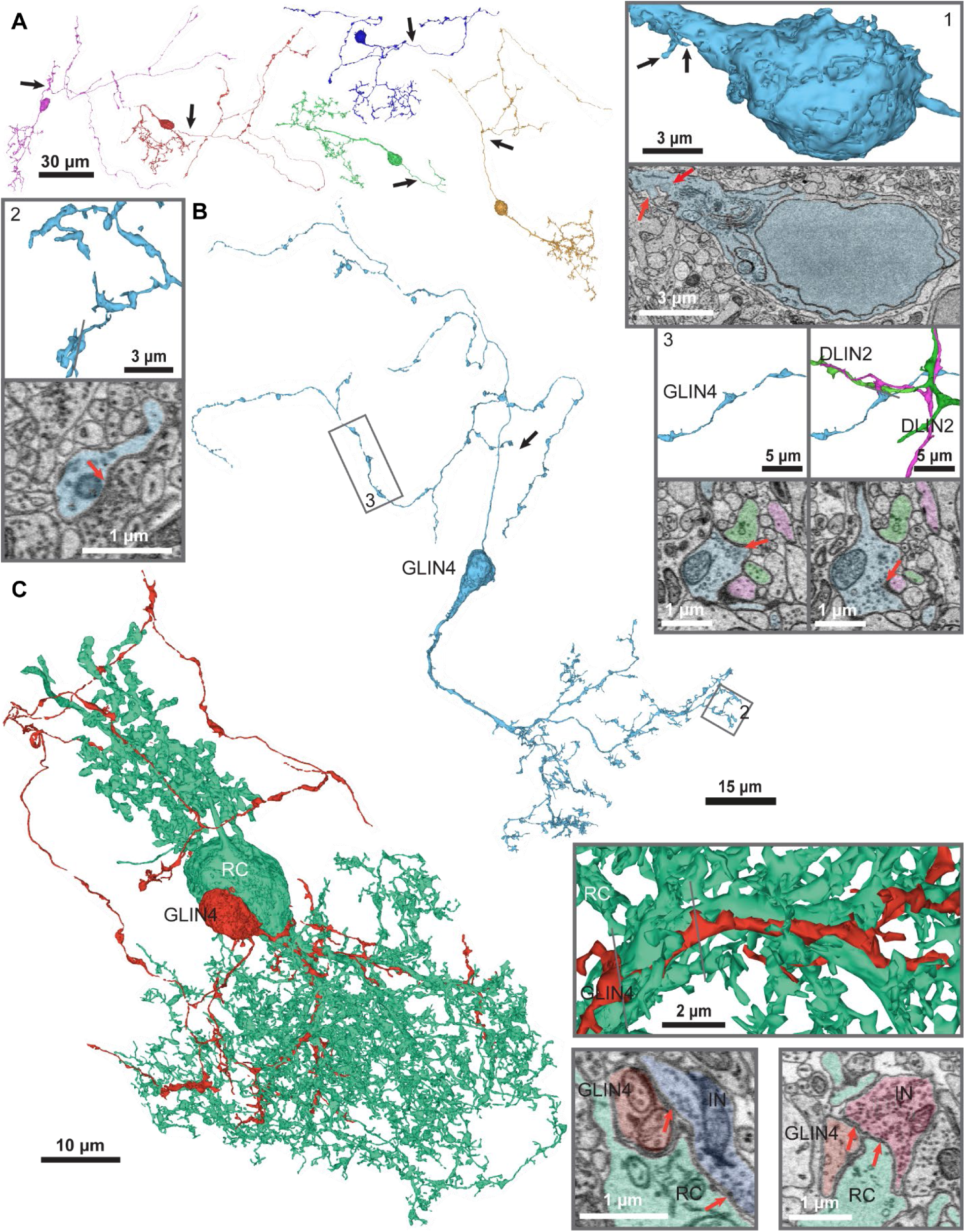
GLIN4. A. Five GLIN4s (arrows: axons). B. Representative GLIN4 with an ovoid nucleus embedded in sparse cytosol, organelles at the base of the primary dendrite, and two short primary cilia (1). The primary dendrite branches into a bushy arbor and axons remain local, branching extensively. (2) Distal dendrite with an input synapse. (3) Axon with amorphous varicosities and synaptic connections onto dendrites two other DLINs (green, magenta). Bottom: cross sections showing synapses (arrows). C. Representative anatomical association of a GLIN4 with a RC. Insets show intertwined dendrites (top) and shared input from another IN (bottom).

*Interactions with PNs.* Synaptic connectivity between GLIN4s and PNs was examined between pairs of GLIN4s and PNs innervating common glomeruli. Among 330 GLIN4-MC pairs, four MC-to-GLIN4 connections and two GLIN4-to-MC connections were found. The inspection of 56 GLIN4-RC pairs yielded two pairs with synapses from the RC ruff onto a GLIN4 axon. One of these connections was reciprocal. Ten GLIN4s had dendrites that entwined dendrites of RCs and shared synaptic input (Fig. 7B).

#### Plexiform layer interneurons (PLINs)

##### PLIN1

*Morphology.* The axonless PLIN1s (n = 6, [view](https://neuroglancer-demo.appspot.com/#!gs://adultob-public/states/PLIN1.json)) had small somata (ø: 7 - 9 µm, *x̅* ± σ: 7.8 ± 0.9 µm) in the PL or GCL. One or two primary dendrites projected radially into the PL and formed bushy ramifications with a diameter of 20 – 30 µm (Fig. 8A, B). Small, occasionally spine-like branches were found on the proximal dendrites and soma (Fig. 8B). Distal dendritic ramifications were immersed within one or more ruffs and exhibited amorphous and flattened, sometimes bulky varicosities (Fig. 8B, C). Three PLIN1s extended a distal tip of one dendrite into a glomerulus.

**Figure 8.**
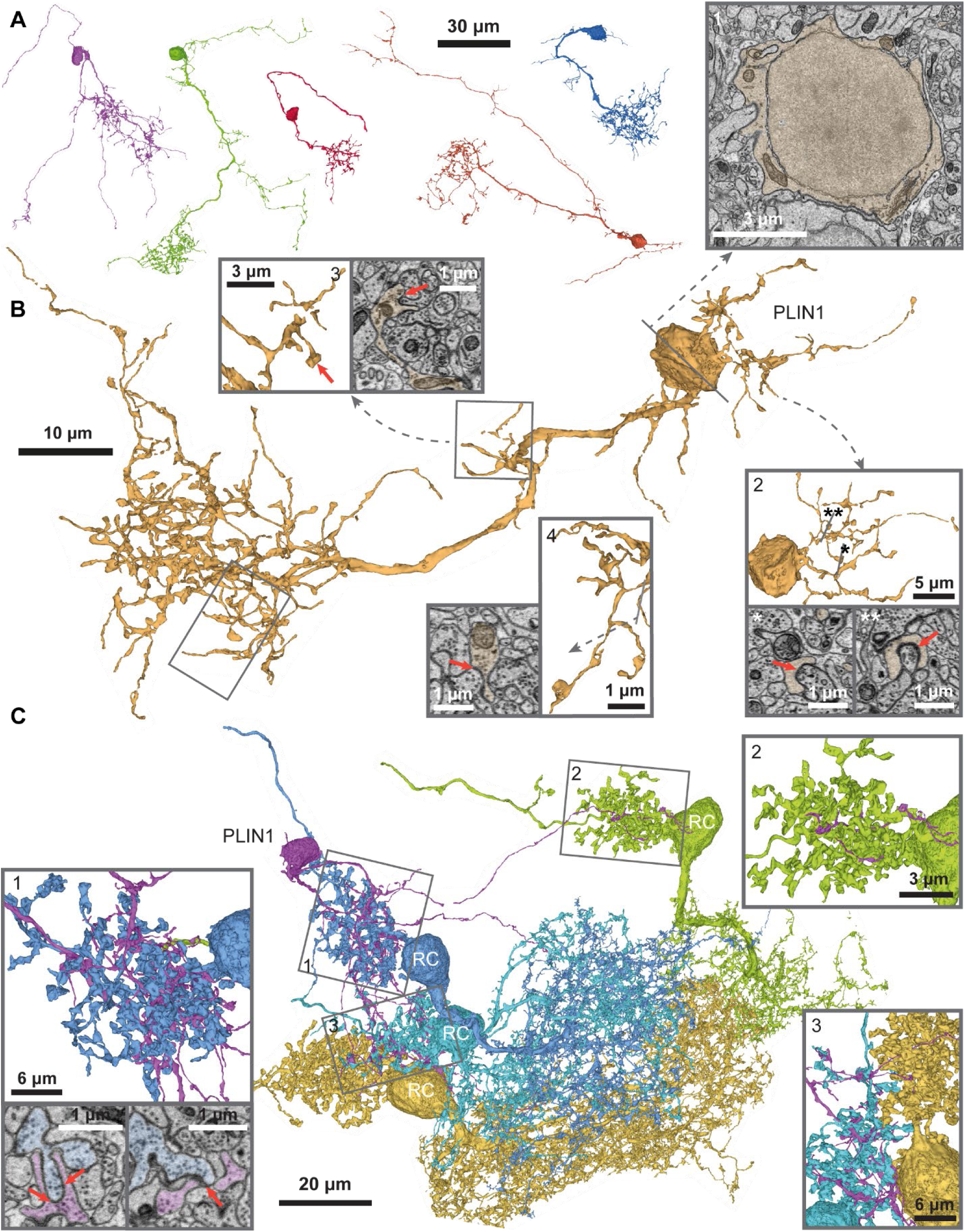
A. Five PLIN1s. B. Another representative PLIN1 with and irregular soma and sparse cytoplasm (1). Proximal dendrites exhibit spine-like protrusions that are exclusively postsynaptic (2; arrows). Distal dendrites have amorphous varicosities with outbound synapses (3, 4; arrows). C. A PLIN1 with distal dendrites immersed into the ruff neuropil of four RCs. Innervation of ruffs can be dense (1) or sparse (2, 3) and contain both inbound and outbound synapses (1, bottom, arrows).

*Interactions with PNs.* While one PLIN1 projecting into GL7 made no connections with PNs, two PLIN1s projecting into GL5 made one or multiple outgoing and reciprocal synapses with one or two out of four MCs. PLIN1s were associated with one to five RCs (1-5, *x̅* ± σ: 3.2 ± 1.3) and 19 of 20 PLIN1-RC pairs were synaptically connected. Connections via the bushy distal dendrites comprised one to five inbound, outbound and/or reciprocal synapses per pair. Out of 19 connections, 13 were reciprocal, four were directed from the RC to the PLIN1, and two were directed from the PLIN1 to the RC.

#### PLIN2

*Morphology.* PLIN2s (n = 11, [view](https://neuroglancer-demo.appspot.com/#!gs://adultob-public/states/PLIN2.json)) were axonless and had small somata that were part of small clusters in the PL (ø: 7 - 9 µm, *x̅* ± σ: 8.0 ± 0.6 µm). A single main dendrite (ø: 0.5 – 1.5 µm, *x̅* ± σ: 0.9 ± 0.3 µm) with terminal ramifications projected into a single glomerulus (Fig. 9 A, B). Four PLIN2s had a second, typically thinner dendritic branch with a length up to ∼30 µm. The main dendrite was studded with appendages at low frequency and regular intervals that could be post-synaptic (Fig. 9B). Appendages varied in length and shape between individual PLIN2s but were consistent within each PLIN2. The tasselated ramifications of the distal dendrites were sparse and restricted to a one glomerulus. In- and outbound, often reciprocal, synapses were located in small varicosities (Fig. 9B,C).

**Figure 9.**
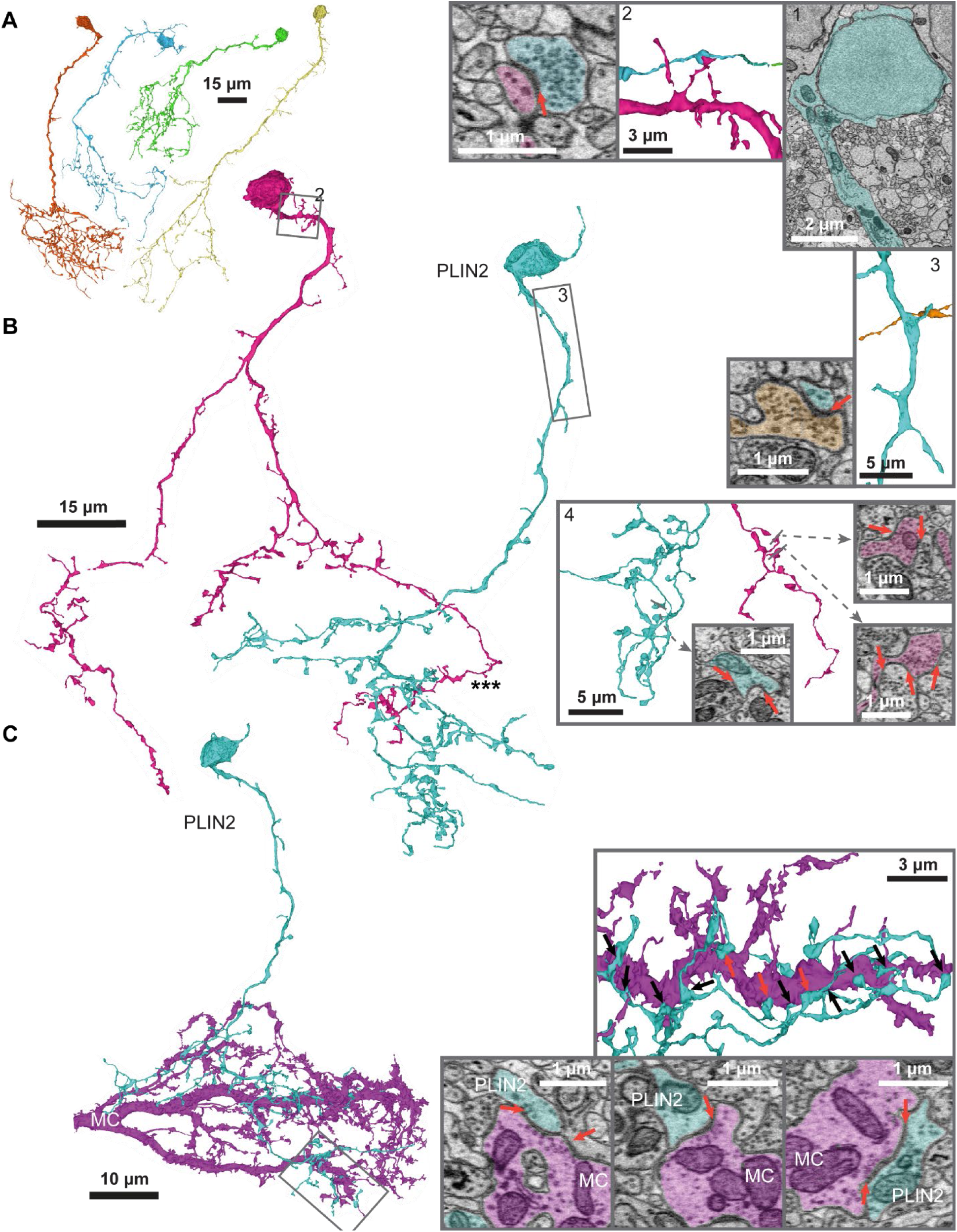
PLIN2. A. Four PLIN2s. B. Two representative PLIN2s with bulbar somata, a thin layer of cytosol, and organelles of the endomembrane system at the base of the primary dendrite (1). Protrusions are sparse along the proximal dendrite and receive synaptic input (2, 3; arrows). Dendrites project toward the GL where they branches into a small, bushy arbor with inbound and outbound synapses on small varicosities (4). C. Example of a PLIN2 connected through a large number of synapses to a MC dendrite in GL5. Insets: top: 12 synapses (arrows) along approximately 20 μm of the MC dendrite. Bottom: Reciprocal (left, right) and unidirectional (center) synapses (red arrows in 1).

*Interactions with PNs.* Synaptic connectivity between PLIN2s and PNs was examined for four PLIN2s associated with GL5, GL6, GL8 or GL9. In GL8, GL6 and GL5, PLIN2s connected to 44 – 60% of the MCs (4/9, 6/12 and 3/5 MCs, respectively). Five pairs were connected by multiple inbound, outbound and reciprocal synapses while the other six pairs were connected by a single reciprocal (3/8), inbound (3/8 or outbound (2/8) synapse. The PLIN2 of GL9 reciprocally connected to two of 13 MCs within a smaller glomerular subcompartment. PLIN2-MC connections could involve up to 25 synapses/pair. None of the nine inspected PLIN2-RC pairs were connected.

##### PLIN3

*Morphology.* PLIN3s (n = 19, [view](https://neuroglancer-demo.appspot.com/#!gs://adultob-public/states/PLIN3.json)) were axonless with mid-sized ovoid somata (ø: 8 - 10 µm, *x̅* ± σ: 9.3 ± 0.7 µm) in the PL and a long primary dendrite extending into the GL (Fig. 10A). Five PLIN3s had an additional small secondary dendrite. Primary dendrites spread widely and branched extensively in the GL, forming a large, often asymmetric, arbor. Proximal dendrites bore spine-like appendages of diverse appearance, most of which received synaptic input (Fig. 10B). Dendrites visited glomeruli (*x̅* ± σ: 3.9 ± 1.7 glomeruli,) where they often ramified and formed amorphous bulgy or flattened varicosities with in- and outbound synapses, primarily with MCs (Fig. 10B, C). PLIN3s may be further subdivided into a varicose and a ribbon-like PLIN3 subgroup (Fig. 10B, C, Supplementary).

**Figure 10.**
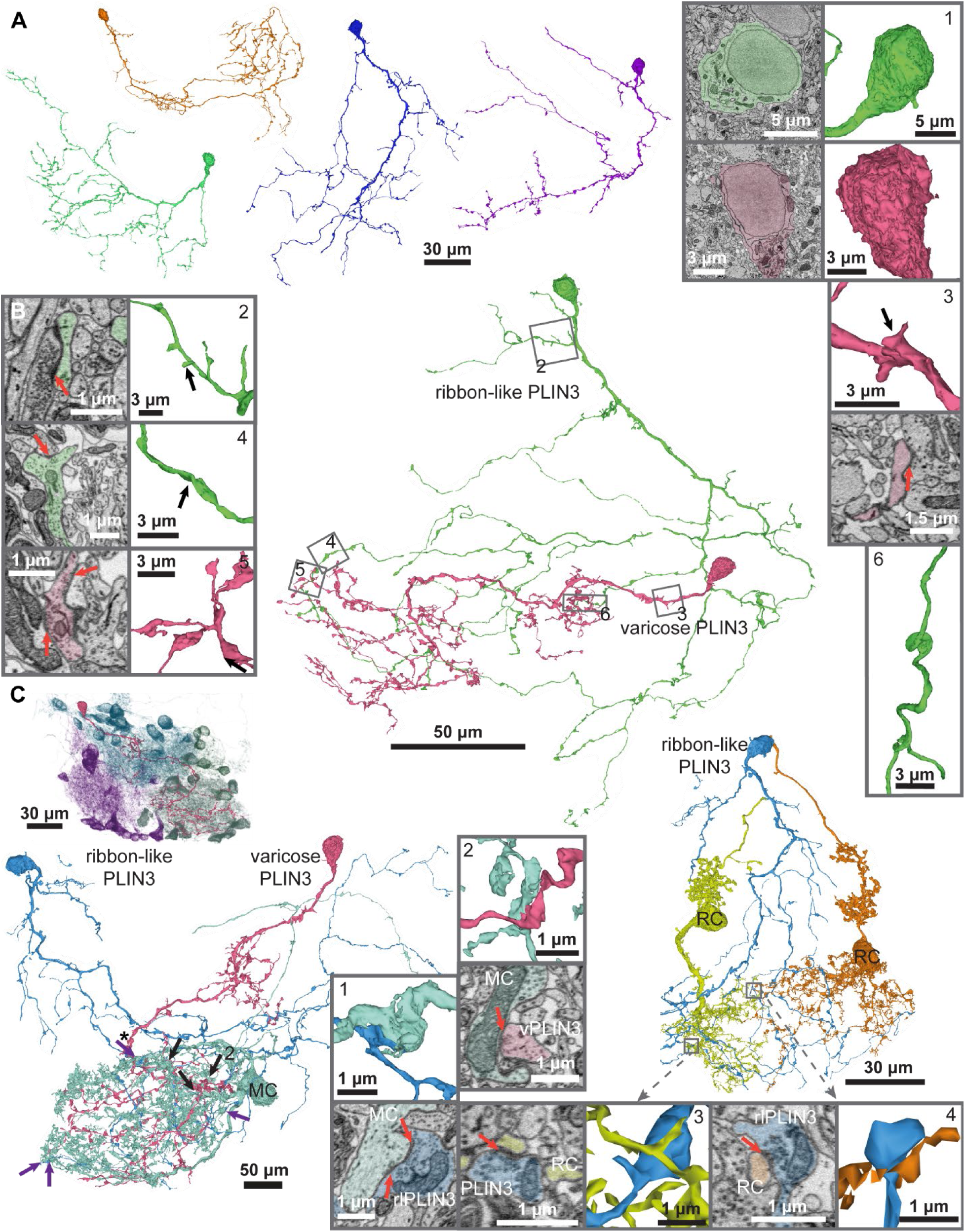
PLIN3. A. Two varicose (green, violet) and two ribbon-like PLIN3s (orange, blue). B. Detailed morphology of a ribbon- like (green) and a varicose (pink) PLIN3. Somata are ovoid, with organelles at the base of the primary dendrite and no protrusions or excrescences. The surface of the ribbon-like PLIN3 is smooth, whereas the varicose PLIN3 is more irregular (1). (2). Proximal branch of the ribbon-like PLIN3 with stubby and long-necked, round-headed spines receiving synaptic input (arrows). (3) Proximal\ branch of the varicose PLIN3 with synaptic input (arrow) onto more irregular, stubby spines. (4, 5) Distal dendrites with varicosities containing inbound and outbound synapses (arrows). Varicosities are flattened in the ribbon-like PLIN3 (4) and bulgy in the varicose PLIN3 (5). (6) Flattened dendrites of the ribbon-like PLIN3 that form ribbon-like curls. C. Connectivity to PNs. PLIN3s are multiglomerular. Top left: association of a representative varicose PLIN3 (red) with GL9 (violet), GL7 (blue), and GL11 (turquoise). Bottom left: A varicose (black arrows) and a ribbon-like (purple arrows) PLIN3 connected to a MC of GL11 via three synapses each. Both PLIN3s form reciprocal synapses (1, 2, arrows). Right: A ribbon-like PLIN3 providing input to the thin dendrites of RCs in GL11 (3, 4, arrows).

*Interactions with PNs.* The connectivity between PLIN3s and PNs was investigated in five varicose and two ribbon-like PLIN3s associated with glomeruli GL4, GL5, GL6, GL7, GL11 and GL12. PLIN3s were strongly connected to MCs: On average 66 % of varicose and 83 % of ribbon-like PLIN3-MC pairs were connected. In both PLIN3 subtypes, most connections were reciprocal (varicose: 47/77 pairs; ribbon-like 17/23 pairs) through reciprocal synapses (60/64 reciprocally connected pairs). 50 reciprocally connected pairs had additional synapses that were outbound (37/50) or inbound (19/50). Among the unidirectionally connected pairs, PLIN3-to-MC connections were more frequent (31/36 pairs) than MC-to-PLIN3 (5/36 pairs). The majority of pairs were connected by two to four unidirectional synapses (range, 1 - 11).

Varicose PLINs connected to RCs in only one of 21 pairs, while ribbon-like PLIN3s connected to RCs in three of five pairs. In all connected pairs, one or two synapses were directed from the PLIN3 onto the RC dendrite.

##### PLIN4

*Morphology.* PLIN4s comprised 33 axonless neurons with flattened or ovoid somata (ø: 8 - 10 µm, *x̅* ± σ: 9.0 ± 0.7 µm) predominantly in the PL, occasionally in the GL or GCL ([view](https://neuroglancer-demo.appspot.com/#!gs://adultob-public/states/PLIN4.json)). PLIN4s had one or two larger (ø: 0.3 - 1.8 µm, *x̅* ± σ: 1.0 ± 0.4 µm) and up to four smaller dendrites (ø: 0.1 - 0.5 µm, *x̅* ± σ: 0.4 ± 0.2 µm) that covered a substantial territory (Fig. 11A). Dendrites were morphologically diverse without a preferred orientation and branched at low to moderate frequency. The overall dendritic appearance was thin and delicate (Fig. 11 B). PLIN4 dendrites projected into the GL and innervated up to five glomeruli (*x̅* ± σ: 2.6 ± 1.2 glomeruli). Dendrites had highly heterogeneous protrusions and appendages (Supplementary). Enlargements of proximal dendrites were usually postsynaptic whereas distal enlargements contained outbound and reciprocal synapses (Fig. 11B, C).

**Figure 11.**
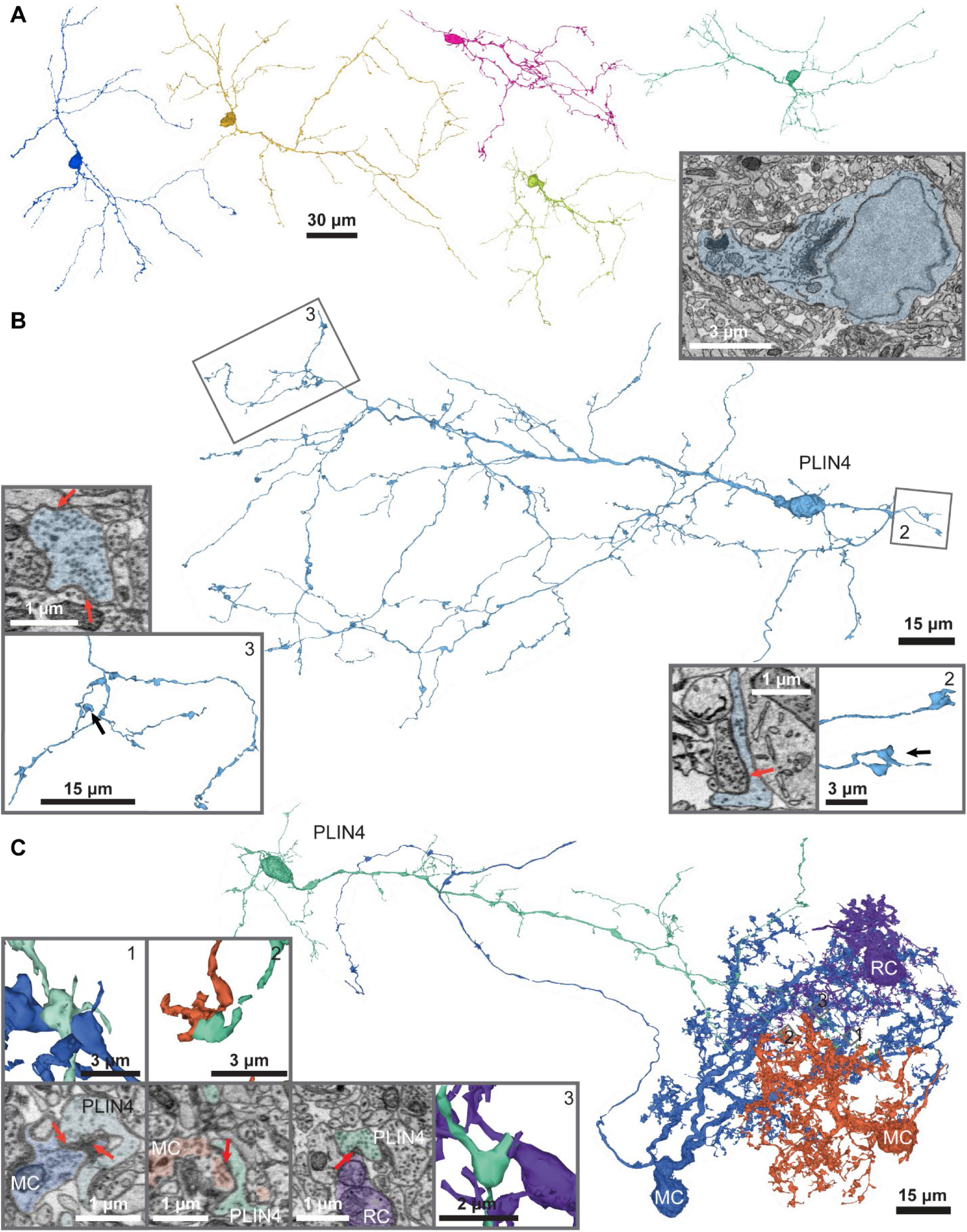
PLIN4. A. Five PLIN4s. B. Representative PLIN4 with an ovoid, uneven soma lacking spine-like protrusions and thin, tortuous dendrites. (1) Nucleus surrounded by sparse mitochondria and ER; most organelles accumulate at the base of the primary dendrite. (2) Long and sparse on the proximal dendrite with a head-like protrusion that contains vesicles and inbound but no outbound synapses (arrow). (3) Bulbous distal varicosities with multiple outbound synapses (red arrows), one of which (black arrow) is shown in cross-section (top). C. Example of a PLIN4 connected to mitral cells and RCs of different glomeruli. Insets show (1) a reciprocal synapse with a MC (arrows), (2) a unidirectional synapse from the PLIN4 onto a MC (arrow), and (3) a synapse from the PLIN4 onto a RC dendrite (arrow).

*Interactions with PNs.* The connectivity between PLIN4 and PNs was investigated for six PLIN4s in glomeruli GL1, GL4, GL6, GL7, GL11 and GL12. PLIN4s connected, on average, 30 % of the MCs (58/196 pairs). Most pairs were reciprocally connected (26/58) or the PLIN4 was presynaptic to the MC (22/58). Unidirectional connections from a MC to a PLIN4 were found in 10/58 pairs. Most connections were mediated by one or two synapses (50/58 pairs) and all reciprocal connections involved reciprocal synapses. PLIN4s connected to 42 % of RCs (15/36 pairs). All connections contained unidirectional synapses from PLIN4 to the RC dendrite. Only one unidirectional synapse was found in the opposite direction. Reciprocal synapses between PLIN4s and RCs were not detected within the inspected glomeruli but observed elsewhere in the dataset (Supplementary).

#### Deep Layer Interneurons (DLINs)

All DLINs were axonless, had a soma in the GCL, and projected dendrites into the GL that typically visited multiple glomeruli. Three morphological classes of DLINs were distinguished based primarily on ultrastructural features, particularly spine-like appendages on proximal dendrites, varicosities of distal dendrites, and the subcellular localization of synaptic connections with PNs. The 61 DLIN1s were characterized by a round, chamfered appearance of their spine-like appendages and flattened, stretched distal varicosities (Fig. 12A). The spine-like appendages of the 81 DLIN2s had a frayed aspect and their distal varicosities were often voluminous and bulky (Fig. 12B). DLIN1s and DLIN2s were further subdivided into three subclasses each. The 38 DLIN3s had proximal dendrites with many long and branched spine-like appendages, resulting in a hairy appearance (Fig. 12C).

**Figure 12.**
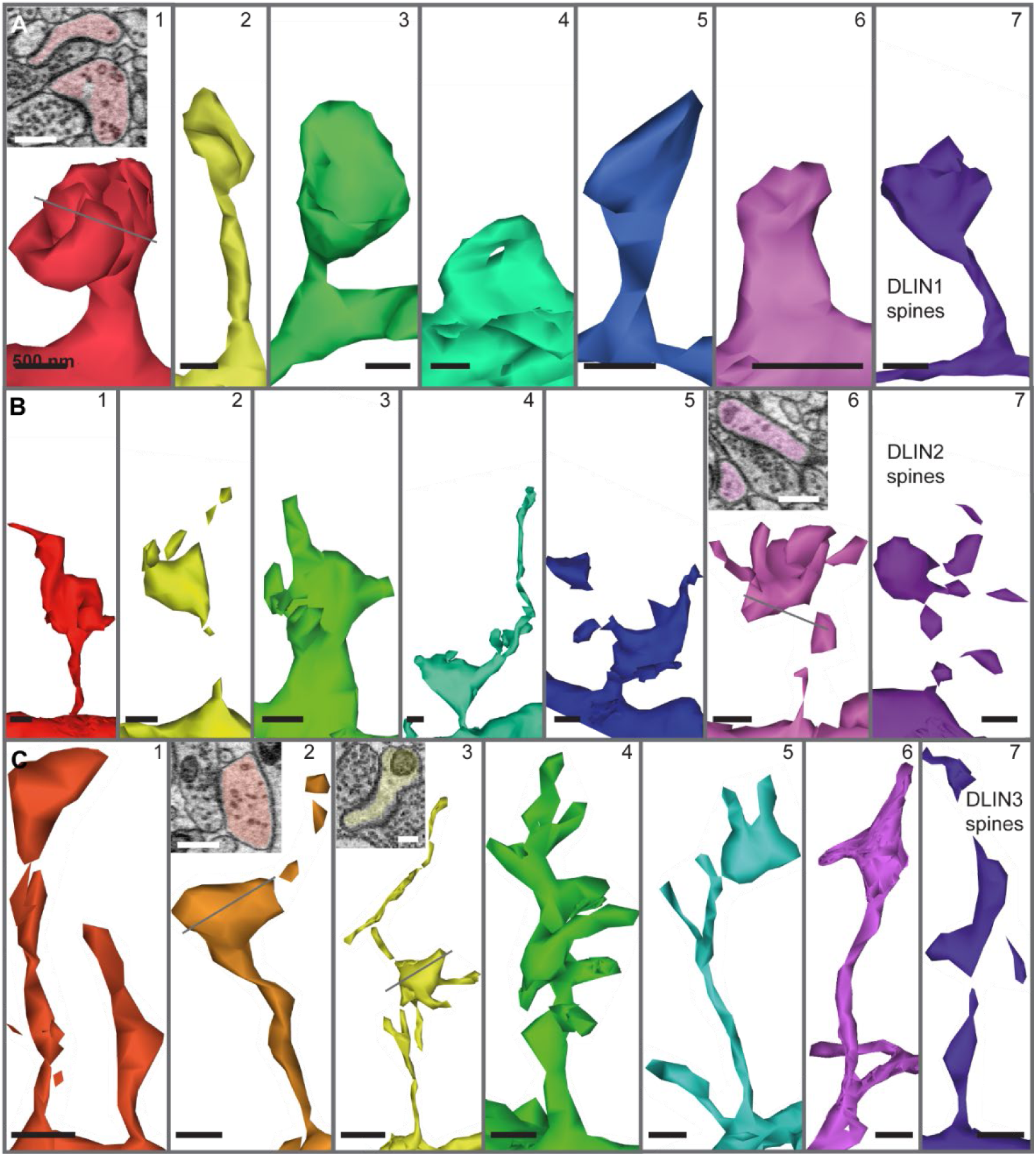
DLIN spines. Spine-like appendages are important features used for the classification of DLINs. Spines receive at least one and often multiple synaptic inputs (indicated by arrows in A.1, B.6, C.2, C.3). A. Spines of DLIN1s. Heads are predominantly rounded. Spines can be short-necked with complex heads (1), cupped (3), or voluminous (5) heads, or long-necked with flattened (2) or voluminous heads (7). Sessile spines can occur either as flattened membrane protrusions (4) or voluminous stumps (6). B. Spines of DLIN2. Spines are typically bulgy and frayed. Necks are of moderate length and typically terminate in bulky, amorphous heads with one or more frayed or pointed protrusions. Some branched (7) and sessile spines (3) are observed. C. Spines of DLIN3s. Spines are typically thin, elongated protrusions several micrometers in length that occur at high frequency. Filopodial as well as mushroom-like branches can arise either along the middle of a spine neck or at the distal end (4, 5). Many spines also exhibit amorphous dilations along the neck that are typically targeted by synapses and continue into a filopodium (2, 3).

#### DLIN1

*Morphology.* DLIN1s (n = 61, [view](https://neuroglancer-demo.appspot.com/#!gs://adultob-public/states/DLIN1.json)) typically had globular or ovoid somata (ø: 8 - 11 µm, *x̅* ± σ: 9.4 ± 0.4 µm) and two types of dendrites: thin basal dendrites in the GCL and larger main dendrites that projected to the GL and ramified in up to seven glomeruli (*x̅* ± σ: 3.2 ± 1.5 glomeruli). Three subclasses could be distinguished: (1) Rough DLIN1s (n = 20) were large, had one or two main dendrites and were covered by a high density of stubby and short spines on the soma and proximal main dendrites. Basal dendrites were delicate with short-necked spines (Fig. 13A). (2) Long-spiny DLIN1s (n = 22) had spines of different lengths, including long-necked spines, at moderate frequency on all deep dendrites. They were smaller than rough DLIN1s and typically located more superficially (Fig. 13B). (3) Sparsely spiny DLIN1s (n = 19) had somata deep in GCL and gave rise to up to four slender main dendrites with stubby and short spines at low frequency (Fig. 13C).

**Figure 13.**
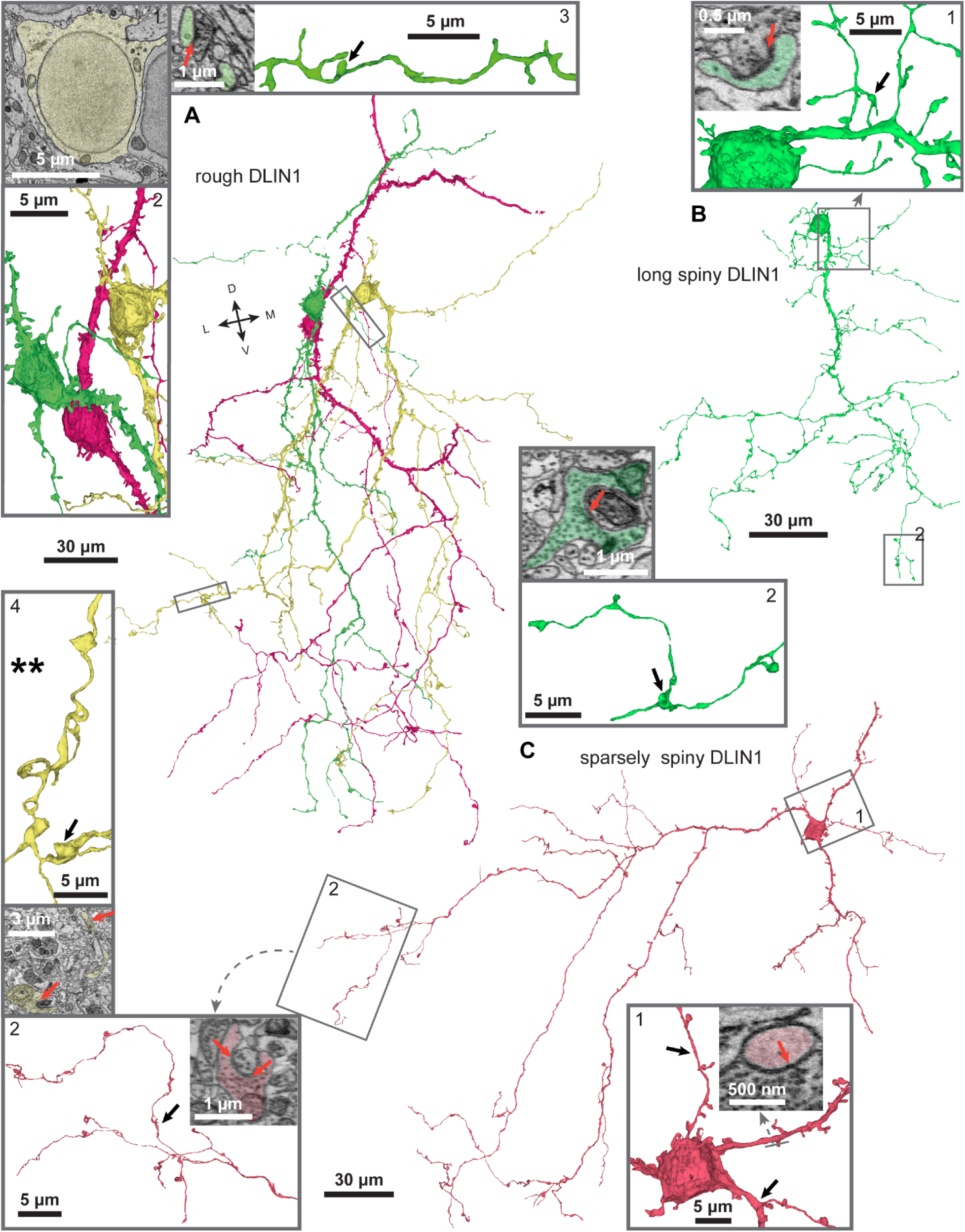
DLIN1 subclasses. A. Three rough DLIN1s (rDLIN1s) with large primary dendrites projecting to the ventrolateral GL. Two rDLIN1s (green, pink) project a secondary large dendrite medio-dorsally, presumably towards glomeruli outside the imaged volume. Within the GL, the widespread distal dendrites bear varicosities at a low to moderate frequency. (1) Somata are ovoid with an egg-shaped nucleus surrounded by moderate amounts of cytosol. (2) Somata and proximal dendrites are densely covered with flattened membrane excrescences and stubby, short-necked, rounded spines. (3) Proximal dendrites are thin and studded with short-necked, round-headed spines at moderate frequency. Inset: Spines receive synapses from thin axons densely packed with small, dark vesicles that originate outside the imaged volume (arrow). (4) Distal dendrite with ribbon-like curls (top). The elongated expansions of DLIN1 dendrites envelop neighboring neurites and contain outbound (bottom, arrows) and inbound synapses (not shown). B. A long-spiny DLIN1 that is characterized by a moderate density of long-necked spines with rounded heads. (1) The soma bears occasional spines but is otherwise smooth. Long-spiny DLIN1s have numerous thin basal dendrites with long- and short-necked spines that are exclusively postsynaptic (inset). (2) Small varicosities of distal dendrites—either bulky or cup-shaped and enveloping adjacent neurites—are sites of synaptic output (top, arrow) and input (not shown). C. A sparsely-spiny DLIN1. (1) The soma bears membrane excrescences and occasional spines. Basal dendrites (arrows), seen in a subset of sDLIN1s, bear predominantly stubby and short-necked spines at low to moderate frequency. Inset: Synapse onto a stubby spine. (2) Varicosities of distal dendrites are mostly flattened, amorphous, or coin-shaped. They are densely filled with synaptic vesicles and contain inbound and outbound synapses (inset, arrows).

*Interactions with PNs.* For two representatives of each subclass the connections to PNs were examined in the glomeruli GL6, GL9, GL11 and GL12. Different DLIN1s innervated different fractions of MCs per glomerulus (rough DLIN1s: 25/85 MCs; long-spiny DLIN1s: 24/53 MCs; sparsely spiny DLIN1s 18/65 MCs). Most connections were reciprocal (rough DLIN1s: 15/25 (60 %), 5/25 (20 %), 5/25 (20 %) reciprocal/outbound/inbound synapses, respectively; long-spiny DLIN1s: 17/24 (71 %), 4/24 (17%), 3/24 (13 %) reciprocal/outbound/inbound synapses, respectively; sparsely spiny DLIN1s: 9/18 (50 %), 8/18 (44 %), 1/18 (6 %) reciprocal/outbound/inbound synapses, respectively). Most connections comprised only one or two synapses. Some synapses from rough DLIN1s onto MCs were very large (Supplementary, Fig. 14B).

Rough and long-spiny DLIN1s connected broadly to the majority of RCs while only one sparsely spiny DLIN1 was connected to a RC (rough DLIN: 7/12 (58%) connected, long-spiny DLIN1s: 7/10 (70%); sparsely spiny DLIN1s: 1/8 (13 %)). Additional synapses from sparsely spiny DLIN1s onto RCs were frequently observed elsewhere in the dataset, suggesting that the connectivity between this DLIN1 subclass and RCs is more complex (Supplementary). In 14 of 15 connections the DLIN1 provided input to the dendrites of the RCs through one to four synapses.

##### DLIN2

*Morphology.* DLIN2s (n = 81, [view](https://neuroglancer-demo.appspot.com/#!gs://adultob-public/states/DLIN2.json)) had ovoid or globular somata (ø: 9 - 12 µm, *x̅* ± σ: 9.9 ± 0.4 µm) that were usually part of a large cluster in the GCL. About half of the DLIN2s gave rise to one or two primary dendrites projecting to the superficial layers and up to three smaller basal dendrites. The remaining DLIN2s had up to four primary dendrites of similar size. DLIN2s were among the largest neurons in the dataset and projected into up to 11 glomeruli (x̅ ± σ: 3.7 ± 1.9, Fig. 16). In contrast to DLIN1s, primary dendrites branched already in PL. Dendrites changed gradually from spiny to varicose along the proximal-distal axis. Presynaptic structures were observed not only within glomeruli but also in the PL (Fig. 15B).

**Figure 14.**
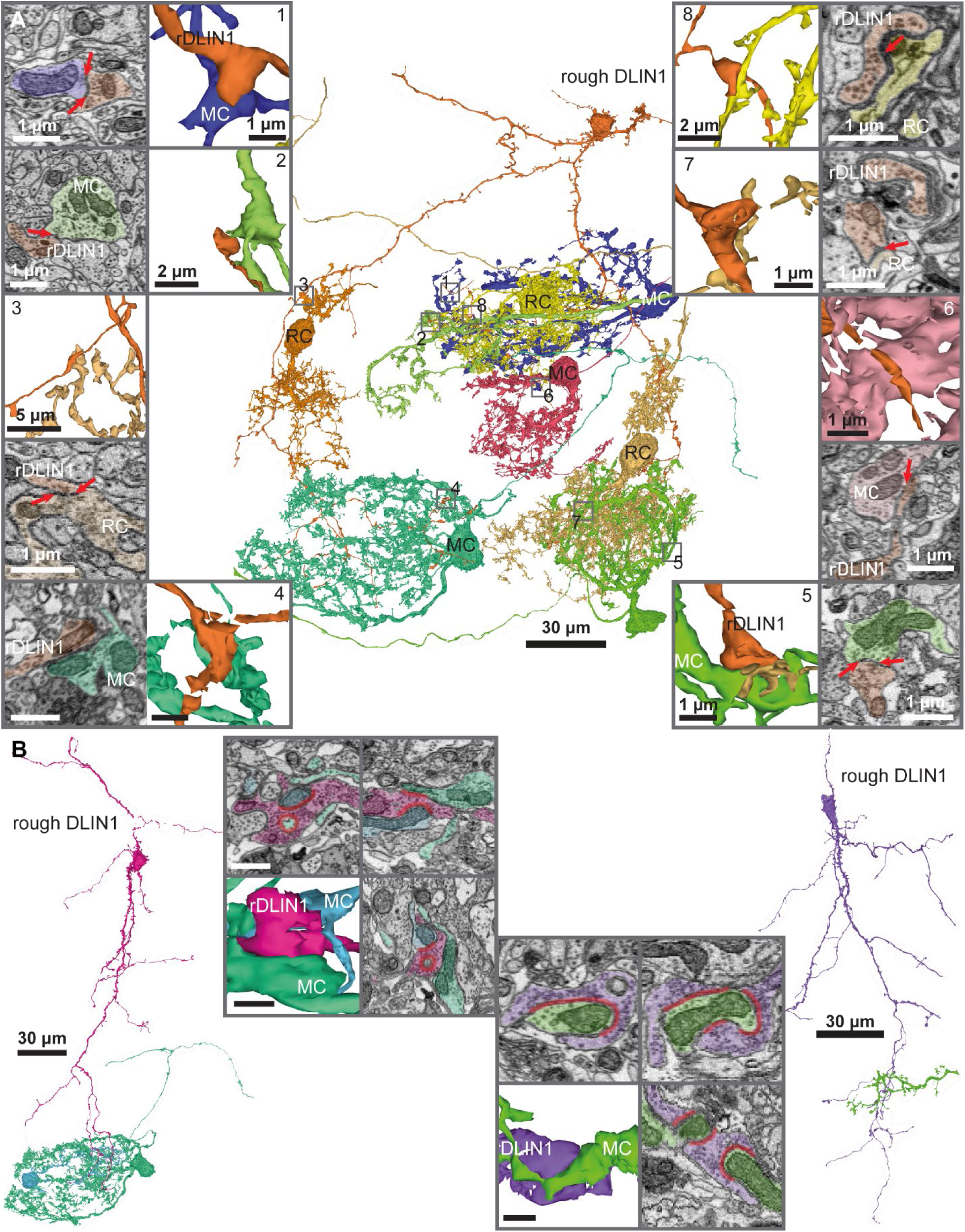
Connectivity between DLIN1s and PNs. A. A rough DLIN1 connecting to MCs and RCs from six different glomeruli. It is reciprocally connected to intermediate (1) and large MCs (2, 4, 5) and one RC (3). In addition, it provides unidirectional input to a small MC (6) and two RCs (7, 8). B. Large synapses between a rough DLIN1 and MCs. The distal dendrite of the rough DLIN1 forms extensive sheaths that wrap around MC dendrites. Synapses onto the MCs can comprise multiple active zones or single zones extending over several hundred nanometers.

**Figure 15.**
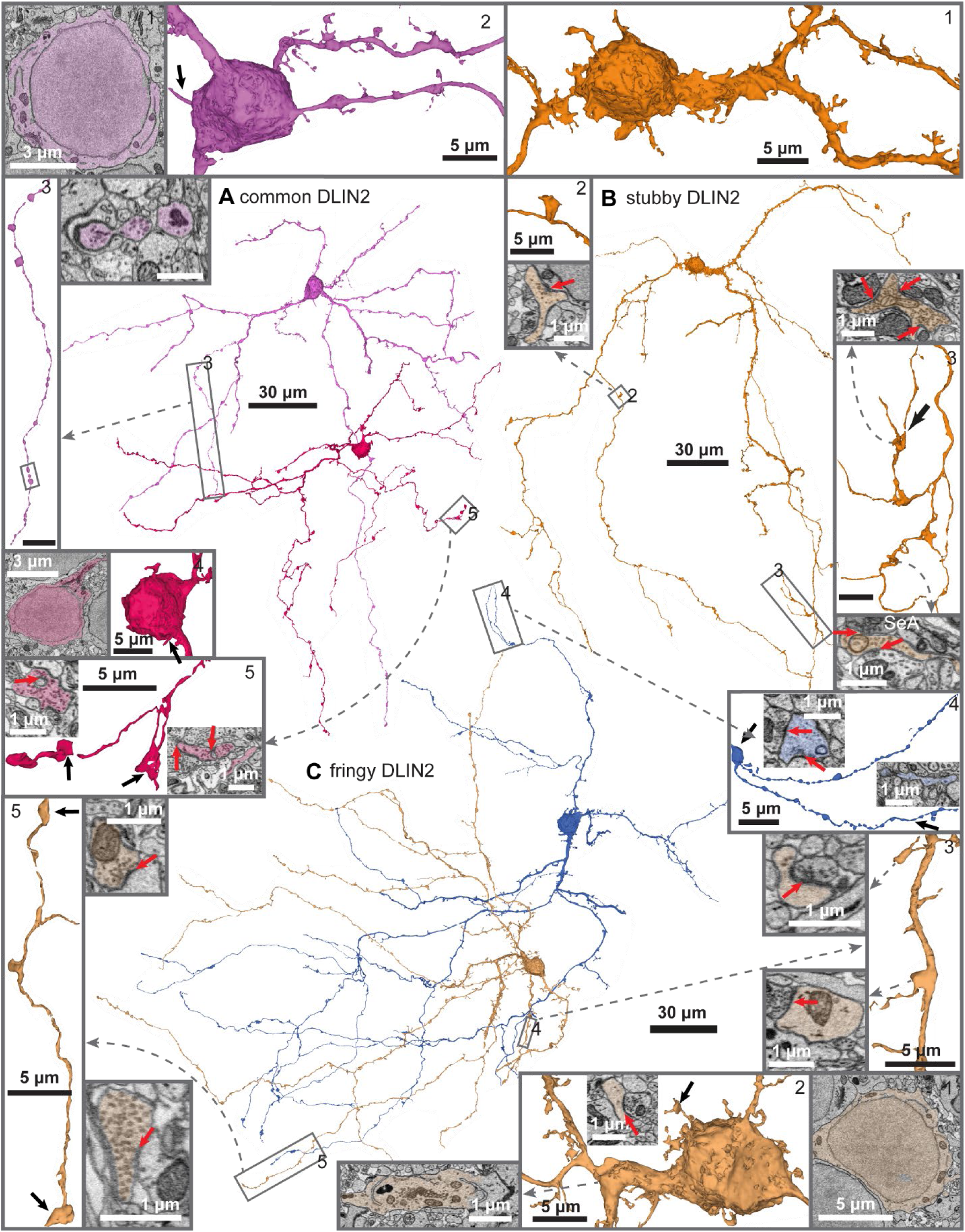
DLIN2 subclasses. A. Two representative common DLIN2s with somata in the superficial and deep GCL. Somata are globular and smooth; spines are rare. The superficial DLIN2 has a smaller soma with less cytosol (1, 4). The primary cilia are among the longest within this class (2, 4, arrows). (2) Proximal dendrites are smooth and exhibit regular bulbar dilations that ear frayed spines. (3) Characteristic beaded dendrites extend through the PL and GCL neuropil with frequent bulbar dilations separated by thin dendritic segments, resulting in a pearl-necklace appearance. These varicosities lack mitochondria but are rich in ER (inset). Pre- and postsynaptic contacts are found at a low frequency (not shown). (5) Varicosities of the distal dendrite are typically voluminous (left) but can also be coin-shaped and contain inbound and outbound synapses (arrows in insets). B. A representative stubby DLIN2s (1) The soma and proximal dendrites are densely covered with irregular membrane excrescences and stubby, short-necked, frayed spines. (2) A varicosity or stubby spine within the PL establishing an outbound synapse onto a soma (arrow). (3) Varicosities of distal dendrites occur at a moderate frequency and are predominantly flattened. Insets, top: Varicosities can form multiple outbound synapses. Bottom: Some varicosities receive input from sensory axons (SeA). C. Two representative fringy DLIN2s. Somata are globular and typically covered with numerous short- and long-necked frayed spines, resulting in a rough appearance (1, 2). Inset: The Golgi apparatus extends deep into the primary dendrite. (3) Spines on proximal dendrites receive synaptic inputs from varicosities of thin neurites packed with small vesicles. (4) Inbound and outbound synapses on a beaded dendrite passing through the PL (insets, arrows). (5) Voluminous varicosities of a distal dendrite in the GL establish synapses onto local neurons (insets).

**Figure 16.**
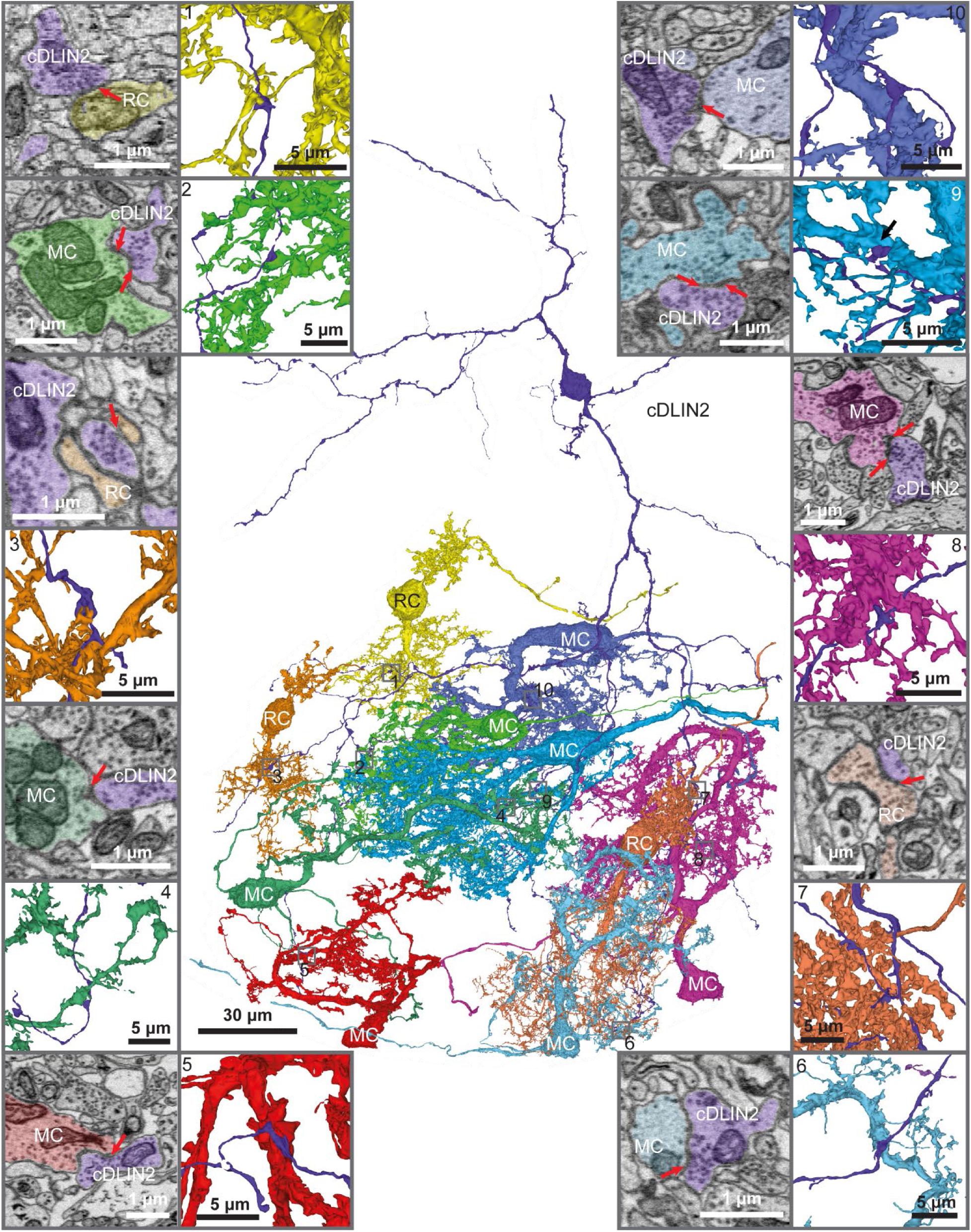
Connectivity between DLIN2s and PNs. A common DLIN2 (cDLIN2) connecting to 19 MCs and nine RCs across nine glomeruli. Fifteen of the connections with MCs (e.g., 2, 8, 9) and two of the connections with RCs are reciprocal (not shown). Unidirectional connections comprised outbound connections to three MCs (e.g., 5, 6, 10) and five RCs (e.g., 1, 3) and inbound connections from one MC (4) and two RCs (e.g., 7). Most connections consist of a single or occasionally two synapses.

DLIN2s were further subdivided into common DLIN2s (n = 52), fringy DLIN2s (n = 22), and stubby- spined DLIN2s (n = 7). Common DLIN2s had smooth somata and carried predominantly short-necked, frayed spines on their dendrites at low to moderate density (Fig. 15A). Fringy DLIN2s had a rugose soma with filopodial protrusions and spines and a spiny proximal dendrite. Both somatic and dendritic spines were typically long-necked, branched and frayed (Fig. 15B). Stubby-spined DLINs were studded with short-necked spines at high density, particularly on the soma and proximal dendrites (Fig. 15C).

*Interactions with PNs.* The connectivity between two common DLIN2s, two fringy DLIN2s and one stubby-spined DLINs to PNs was examined in glomeruli GL1, GL3, GL4, GL5, GL6, GL7, GL8, GL9, GL11 and GL12. DLIN2s typically connected selectively to few MCs within each glomerulus (common: 10/12 glomerulus-DLIN2 pairs; fringy: 4/7 glomerulus-DLIN2 pairs; stubby-spined: 3/5 glomerulus- DLIN2 pairs). Nevertheless, all DLIN2s also connected to a substantial fraction (>40%) of all MCs within at least one glomerulus. The average connection rate to MCs was 22% (33/150 pairs), 26% (18/68) and 29% (19/65) for common, fringy and stubby-spined DLIN2s, respectively. Most connections of common and fringy DLIN2s were mediated by reciprocal synapses (25/33 and 13/17, respectively). DLIN2-to-MC connections (6/33 and 3/17, respectively) and MC-to-DLIN2 connections (2/33 and 1/17, respectively) were less frequent. In stubby-spined DLIN2s, outbound (10/17) and reciprocal (7/17) connections but no unidirectional inbound connections were found. The number of synapses per connection was low for all DLIN2 subclasses, with only eight of 69 pairs being connected by three or more synapses.

Common DLIN2s connected to nine of 33 RCs (27 %). Two connections (20 %) were reciprocal, six connections (60 %) were outbound onto the RC dendrite, and two connections (20 %) were inbound. Outbound synaptic connections onto the RC dendrite were found in 12 (17 %) stubby-spined DLIN2- RC pairs. None of the 14 fringy DLIN2 were connected to RCs.

##### DLIN3

*Morphology.* DLIN3s (n = 38, [view](https://neuroglancer-demo.appspot.com/#!gs://adultob-public/states/DLIN3.json)) had a hairy appearance because all neurites in the deeper layers were densely covered with thin, long (>2 μm) protrusions (Fig. 17A). Most somata (ø: 8 – 10 μm, *xxx* ± σ: 8.9 ± 0.3 μm) were located in a large cluster within the GCL. DLIN3s gave rise to one or two primary dendrites and up to three fine, spiny basal dendrites. Spine-like appendages often received more than one inbound synapse and could also make outbound synapses (Fig. 17A). At the distal dendrites in and outbound synapses were found in compact, bulky varicosities associated with pointy tips (Fig. 17A).

**Figure 17.**
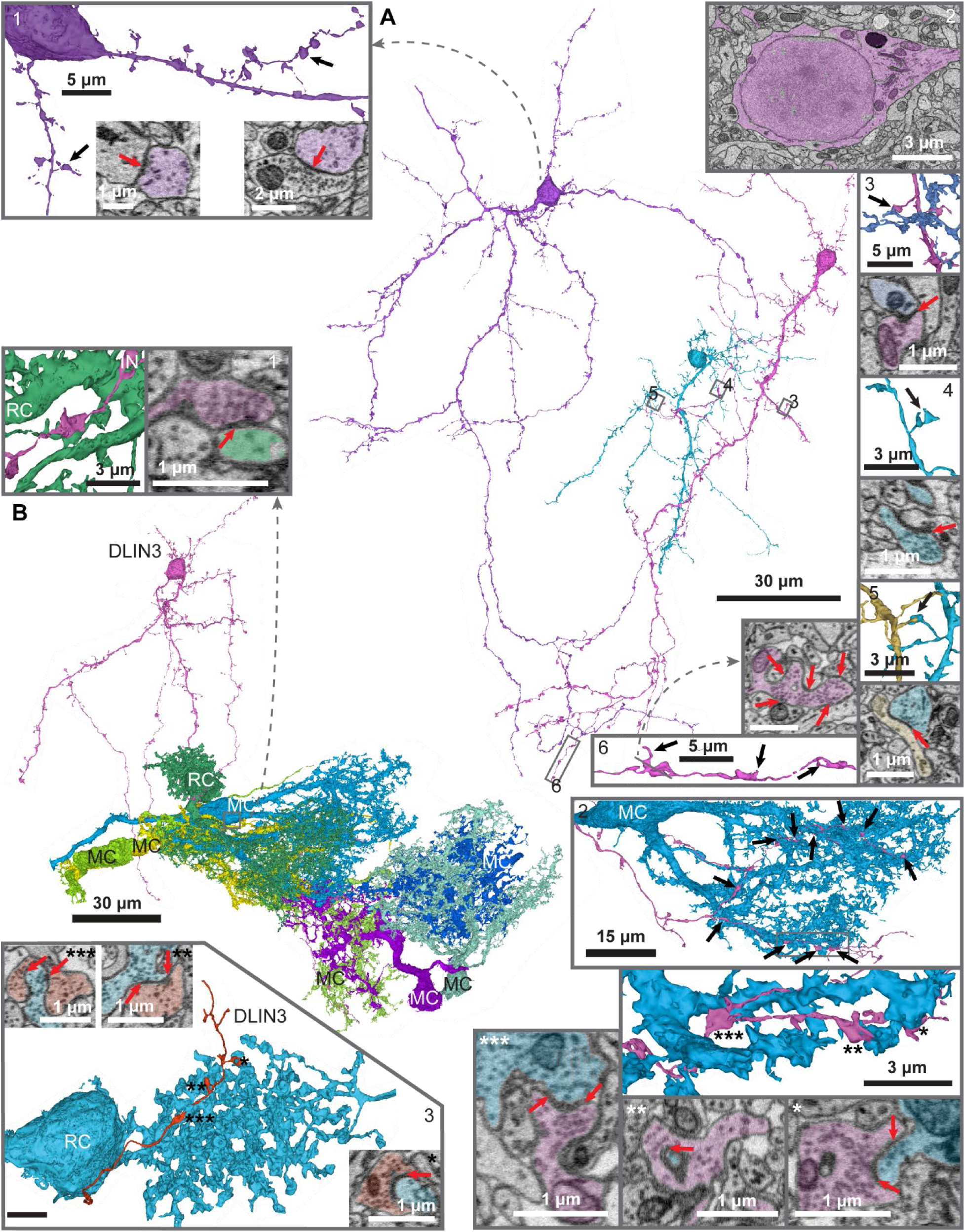
DLIN3. A. Three DLIN3s with typical ovoid to globular, smooth somata occasionally bearing long spines. (1) Hairy spines on proximal dendrites. Some spines contain large vesicular structures resembling synaptic specializations (left; arrow). Right: A spine receiving synaptic input (arrow). (2) Soma with a thin layer of cytosol surrounding the nucleus and a concentration of ER, Golgi apparatus, and mitochondria at the base of the primary dendrite. (3–5) Details of spines within the GCL and PL with outbound synapses (arrows) targeting neurites of presumed other DLINs. (6) Compact, voluminous varicosities in the distal dendrite with pointed tips (black arrows). The inset (top) shows multiple outbound synapses (arrows). B. A DLIN3s connected to seven MCs and one RC across three glomeruli. Connections comprise five reciprocal DLIN3-MC connection, one unidirectional DLIN3-to-MC connection, and one unidirectional DLIN3-to-RC connection. (1) The connection onto the RC consists of a single synapse onto the RC dendrite (arrow). (2) The DLIN3 forms 11 synapses across the dendrite of a single MC (arrows). Center: Three dendritic varicosities closely enveloping the MC dendritic shaft. Bottom: Reciprocal (*, ***) and unidirectional (**) IN-to-MC synapses (arrows). (3) Three reciprocal synapses between another DLIN3 and a RC at the ruff (inset, arrows).

*Interactions with PNs.* Six DLIN3s were investigated for their connectivity to PNs in glomeruli GL7, GL9, GL11 and GL12. Connections were found in 41 of 144 possible pairs with reciprocal connections dominating (30/41), typically through reciprocal synapses (28/30). Additional outbound and inbound synapses were found in nine and six pairs, respectively (1 – 11 synapses/pair; *xxx* ± σ: 3.3 ± 3.0 synapses/pair, Fig. 17B). In addition, DLIN3s were exclusively presynaptic to seven MCs and postsynaptic to four MCs. These connections were mediated by one or two synapses. Three connections were detected between DLIN3s and RCs. These connections included reciprocal synapses at the ruff and outbound synapses onto the RC dendrite (Fig. 17B).

#### Validation of cell type classification

The anatomical and ultrastructural classification of neuron types was performed by a human expert. To validate the result, the same neurons were independently classified by four additional neuroscientists. Neurons were pre-grouped into PNs and INs, and neuroscientists were asked to assign all neurons to a maximum of 20 classes. No further constraints were provided to define neuron classes. To evaluate the consensus, we represented neurons as nodes in an undirected graph where edge weights represented the number of neuroscientists who assigned a given neuron pair to the same subgroup. The resulting adjacency matrix showed obvious clusters, indicating substantial consensus (Fig. 18A). Consensus was particularly strong among PN types (MCs and RCs) and small IN types (most GLINs and PLINs). We therefore conclude that our classification of neuron types captures systematic anatomical features of distinct classes of neurons. Anatomical information informative for classification is summarized in Table 3.

**Figure 18.**
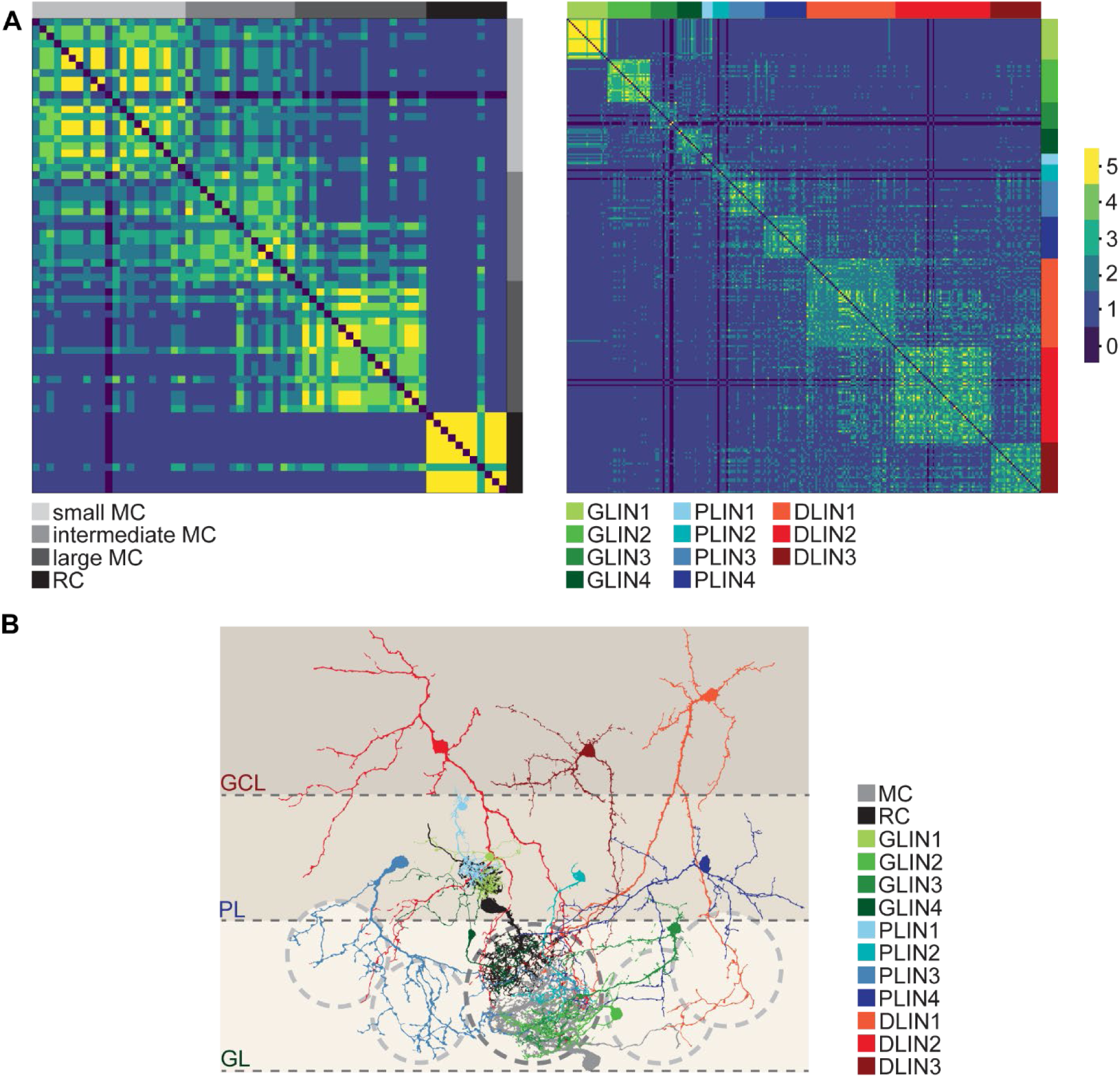
Classification of neuron types: validation and summary. A. Validation of classifications. The assignment of cells into distinct classes by five independent experts is represented as an adjacency matrix of the joint classification graph for PNs (left) and INs (right). B. Association of neuron types with a glomerulus, schematically illustrated by overlays of reconstructed neurons representing different types (color-coded). Dark dashed line approximates the outline of a glomerulus defined by the dendritic arbors of MCs and RCs. Light dashed lines represent adjacent glomeruli. The GL contains the monoglomerular GLIN1–GLIN4. The PL features four predominantly multiglomerular IN types (PLIN1–PLIN4). The GCL contains three main classes of INs (DLIN1–DLIN3) that are further divided into subclasses.

#### Sensory input to OB neurons

All principal neurons (MCs and RCs) received pronounced input from axons of sensory neurons, which can be identified by their characteristic dark cytoplasm (Pinching and Powell, 1971b). Sensory input to IN dendrites, in contrast, was rare. To characterize sensory input to INs more systematically we examined the presence of synapses between sensory axons and 2 – 11 representatives of each IN subclass (Table 4). Most GLINs and PLINs (GLIN1, GLIN4, PLIN1, PLIN2, varicose PLIN3, PLIN4) received no direct input from sensory axons. Sensory input was detected in at least one GLIN2, GLIN3 and ribbon-like PLIN3. These neurons received at most one sensory synapse, some of which were small. In DLINs, particularly DLIN1s, sensory inputs were more frequently detected, with a maximum of 13 sensory synapses per neuron (Fig. 19A, Table 4). Hence, deep INs received more sensory input than superficial INs but sensory input to INs was generally rare. We further observed that some INs, most of which were identified as DLINs, made reciprocal synapses with boutons of sensory axons. In some cases, INs also made output synapses onto sensory axons (Fig. 19B). These synaptic contacts were not observed in the vicinity of output synapses of sensory axons onto MCs.

**Figure 19.**
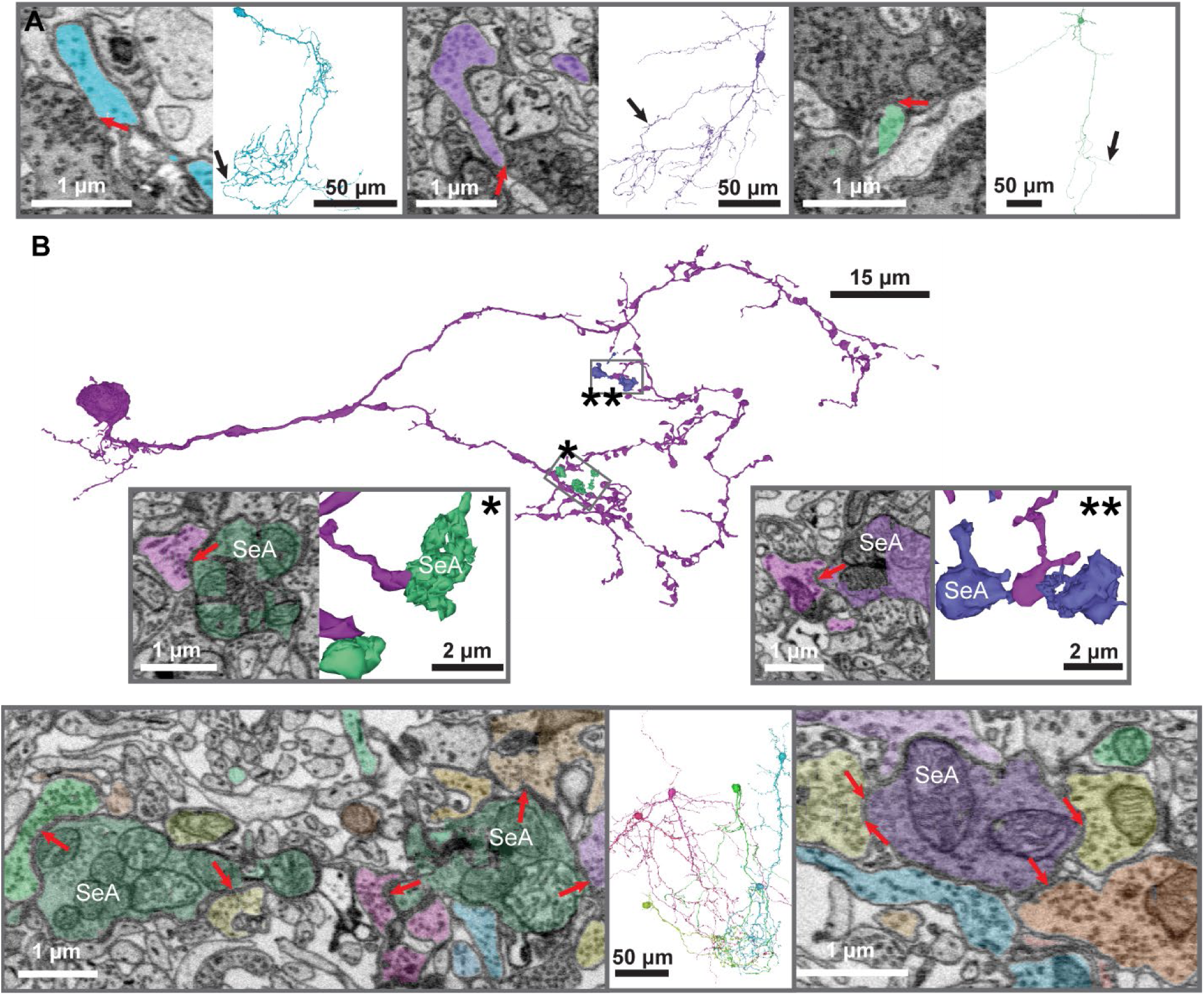
Connectivity between INs and sensory axons. A. Synapses of sensory axons onto a PLIN3 (left), a DLIN3 (center) and an fDLIN2 (right). B. Synapses (arrows) onto sensory axons (SeA) from a GLIN3 (top) and synaptic contacts between multiple other INs and SeAs (bottom). All highlighted neurites provide input to the SeAs (some synapses are out of the focal plane). Synapses onto the SeA frequently target varicosities and can be reciprocal. Bottom center: Six fully reconstructed neurons providing input onto SeAs. Right: Synapses between INs and a SeA; a reciprocal synapse is visible on the left.

#### Neuronal circuit organization

Anatomical features of neuron types in the zebrafish OB and their associations with glomeruli are summarized in Fig. 18B. Along the radial axis two trends could be observed (Fig. 18B, Fig. 20A): First, GLINs and superficial PLINs were monoglomerular, deeper PLINs were multiglomerular, and DLINs innervated the highest number of glomeruli. Hence, the number of innervated glomeruli increased with depth. (2) Within a glomerulus, superficial INs (GLINs, PLIN1-3s) connected to most PNs while the fraction of innervated PNs was variable in deeper INs (PLIN4s, DLINs). Hence, intraglomerular connectivity tends to become sparser and potentially more selective with increasing soma depth.

**Figure 20.**
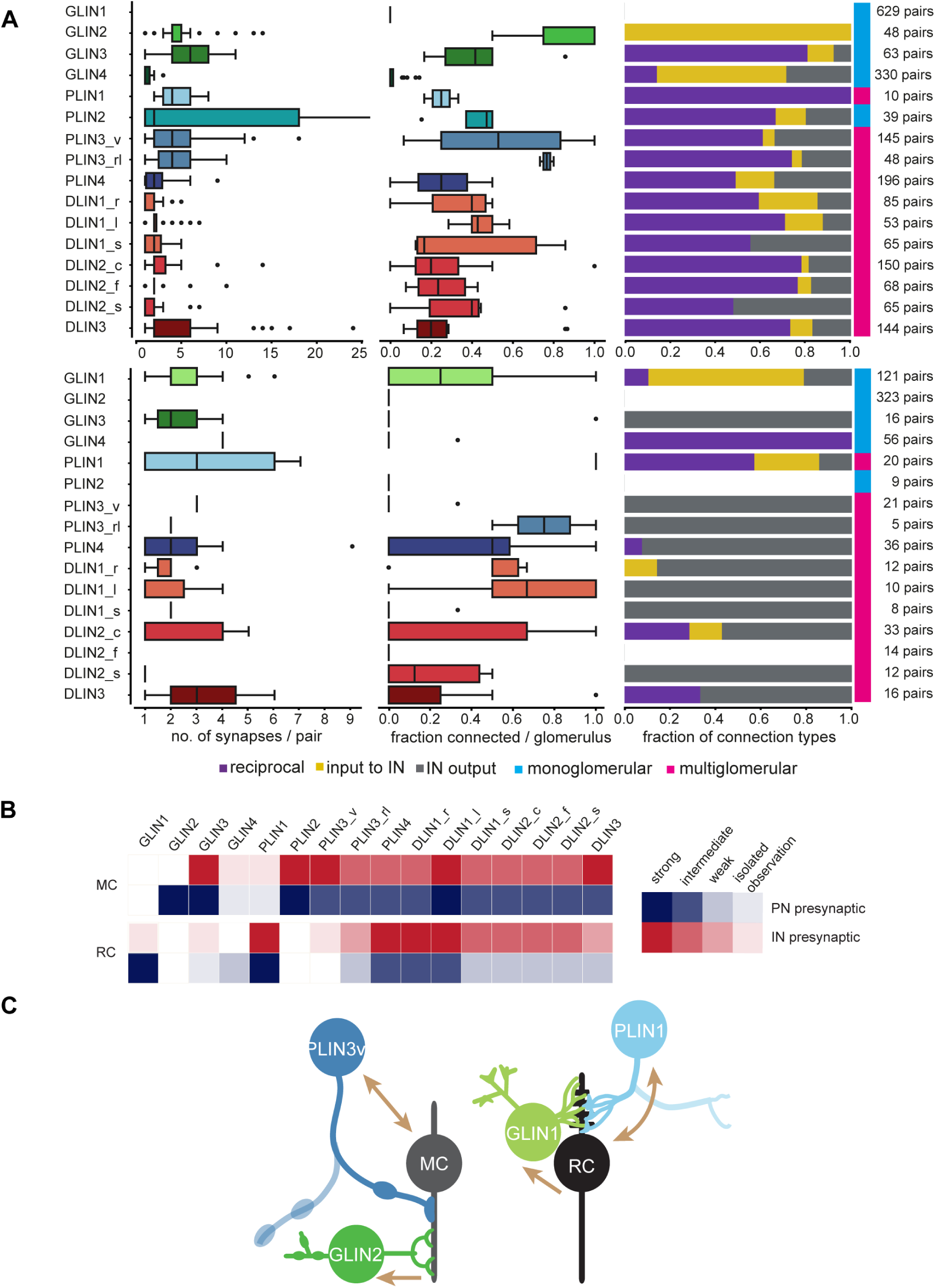
Intraglomerular connectivity: summary. A. Summary of the connectivity between IN and PN classes in the glomerular neuropil. Top: IN-MC connectivity. Bottom: IN-RC connectivity. B. Qualitative summary of observed connection strengths between IN classes and MCs or RCs, respectively. This qualitative assessment of connection strength is based on the intraglomerular connectivity shown in (A) and additional non-systematic observations of connections outside the glomerular neuropil. C. Two\ synaptic microcircuits anchored to MC and RCs. Microcircuits share a similar architecture but are anatomically distinct. Monoglomerular GLINs (GLIN2s and GLIN1s) receive unidirectional input from MCs and RCs, respectively, and project to extraglomerular targets. Multiglomerular PLINs (varicose PLIN3s and PLIN1s) are reciprocally connected to MCs and RCs, respectively. These microcircuits may mediate lateral inhibition and recurrent inhibition (gain control) in subnetworks involving different types of output hneurons.

The annotation of intraglomerular synapses in representative INs revealed specific connectivity between neuron types. While DLINs made synapses with both MCs and RCs, GLINs and most PLINs were primarily or exclusively connected to a single PN type. GLIN2s, GLIN3s, PLIN2s and varicose PLIN3s were primarily connected with MCs. While GLIN2s were exclusively postsynaptic to MCs, GLIN3s, PLIN2s and varicose PLIN3s were reciprocally connected. GLIN1s and PLIN1s, in contrast, were primarily or exclusively associated with RCs. Connections between RCs and PLIN1s were bidirectional while connections to GLIN1s were exclusively outgoing. Hence, MCs and RCs are selectively connected to distinct subsets of GLINs and PLINs, indicating that these PN types are part of different subnetworks (Fig. 20C) containing similar connectivity motifs. In both subnetworks, distinct multiglomerular PLINs (varicose PLIN3 and PLIN1) were reciprocally connected to PNs whereas specific GLINs (GLIN2 and GLIN1) received unidirectional input from principal neurons. Both GLIN types made very few synapses onto principal neurons and had axons that terminated on extraglomerular targets. These results indicate that microcircuits within MC and RC subnetworks are distinct but share similar architectures (Fig. 20C).

The comprehensive volumetric reconstruction of OB neurons revealed also specific non-synaptic associations between neurons. Dendrites of MCs and RCs were often directly adjacent and followed each other over large distances (micrometers), often without making obvious synaptic contacts. Hence, MCs and RCs formed pairs that were tightly linked anatomically whereas synaptic connections were sparse (Fig. 3). Similarly, the entire dendrite of GLIN1s was tightly enveloped in ruffs but received little synaptic input from the RC (Fig. 4B). The purely postsynaptic dendrites of GLIN4s and RCs showed a similar tight anatomical association without chemical synapses (Fig. 7). These anatomical associations may support non-synaptic communication between cells, for example, through paracrine messengers.

## Discussion

We present a comprehensive anatomical classification of neurons in the zebrafish OB. Volumetric EM-based reconstructions provided nanometer resolution at whole-neuron scale, allowing us to link ultrastructural features such as spine shape and neurite diameter to neuronal morphology. Subcellular information contributed significantly to the classification of neuron types, for example, by revealing consistent differences in spine morphology between DLIN types. The ability to annotate synapses, along with the reconstruction of many neurons in the same specimen, enabled systematic insights into the connectivity between neuron types. Currently, circuit diagrams of the OB typically represent neuron types and their spatial relations without defining their connectivity (Kermen et al., 2013; Nagayama et al., 2014) because knowledge about synaptic connectivity is limited in any vertebrate species. The large-scale analysis of neuron types at synaptic resolution thus provides novel structural insights into the organization of the OB that is critical to delineate neuronal microcircuits.

### Glomerular organization

One of the hallmarks of the first olfactory processing center in vertebrates and invertebrates is the segregation of sensory inputs into discrete glomerular processing channels. In some areas of the teleost OB including the ventro-lateral OB of zebrafish, however, neuropil in the afferent layer appears diffuse and glomeruli cannot be clearly delineated histologically (Oka et al., 1982; Satou, 1990; Baier and Korsching, 1994; Byrd and Brunjes, 1995a; Braubach et al., 2012a; Braubach and Croll, 2021). Based on light microscopic data it has been suggested that the ventro-lateral OB of zebrafish contains MCs with multiple dendritic tufts that innervate combinations of small glomerular units (“microglomeruli”) (Braubach and Croll, 2021), resembling the organization of the accessory rather than the main olfactory bulb in mammals. However, our comprehensive analysis revealed that both uni- and multidendritic MCs consistently innervate one of multiple defined subvolumes within the neuropil. The ventro-lateral OB is therefore organized into densely packed yet clearly segregated glomeruli, reinforcing the notion that distinct glomerular channels are a canonical feature of the OB. Moreover, these observations confirm that MCs are typically uniglomerular even when they are multidendritic (Byrd and Brunjes, 1995b; Wanner et al., 2016).

### Anatomical classification of neuron types

Previous anatomical, immunohistochemical and functional studies described different neuron types in the zebrafish OB (Friedrich and Laurent, 2001; Edwards and Michel, 2002a; Fuller et al., 2006a; Bundschuh et al., 2012; Zhu et al., 2013; Miyasaka et al., 2014; Wanner et al., 2016) and classic studies in goldfish described cell types based on high-resolution images (Kosaka and Hama, 1979; Kosaka and Hama, 1982b; Kosaka and Hama, 1982a). However, a comprehensive taxonomy of cell types in OB has been lacking, as in most other vertebrates. The rich information provided by hundreds of EM-based reconstructions enabled a deep classification of neuron types that was cross-validated by independent expert annotations. As the zebrafish OB is a model to study odor representations, our systematic description of neuron types may provide an anatomical foundation for experimental and computational studies. Moreover, the comprehensive classification of neurons clarifies relationships between OB cell types across species.

Consistent with previous studies we found a clear distinction between the two major types of PNs, MCs and RCs. It remains, however, unclear how these neuron types may be related to mitral and tufted cells in mammals. While no obvious mammalian equivalent of RCs has been described, some MC subtypes in the teleost OB may correspond to tufted cells. As MCs and tufted cells project to different target areas (Fukunaga et al., 2012; Igarashi et al., 2012), this hypothesis may now be addressed by mapping projections of MC subtypes in zebrafish.

Because MCs make few direct synaptic contacts, information processing in the OB critically depends on polysynaptic interactions mediated by INs. We identified 11 IN types with additional subclasses that shared many similarities with neurons previously described in the OB of other species. Among GLINs and PLINs, we found monoglomerular INs resembling mammalian periglomerular cells (GLIN1-3s, PLIN2s), as well as more extended, oligo-glomerular neurons (PLIN1s, PLIN3s, PLIN4s) that share structural features with short-axon cells (Aungst et al., 2003; Kiyokage et al., 2010). GLIN1s and GLIN2s closely match neuron types referred to as perinest cells and mixed-synapse cells in goldfish (Kosaka and Hama, 1982b). Multiple types of DLINs are likely to correspond to granule cells in other vertebrate classes, as concluded based on their soma location, synaptic connectivity and dendritic ultrastructure.

The mammalian OB contains an excitatory IN type, the external tufted cell, that relays information from sensory neurons to local MCs and potentially modulates sensory signals (Hayar et al., 2004b; Hayar et al., 2004a). A defining feature of external tufted cells is direct sensory input (Hayar et al., 2004b; De Saint Jan et al., 2009), which was not prominent in any IN type in the zebrafish OB. Among PNs, small MCs shared some anatomical features with external tufted cells such as the presence of axon collaterals (Liu and Shipley, 1994) but the resemblance is weak, and no output synapses of small MCs onto other MCs were detected. Hence, our results provide no evidence for an equivalent of the external tufted cell in zebrafish.

In summary, our results support the notion of a broad conservation of neuron types in the OB across vertebrates, with some notable exceptions such as external tufted cells and RCs. Consistent with this interpretation, neurophysiological results indicate a functional conservation of neuronal computations such as pattern decorrelation and divisive normalization in the OB of zebrafish and rodents (Friedrich and Laurent, 2001; Niessing and Friedrich, 2010; Zhu et al., 2013; Gschwend et al., 2015; Roland et al., 2016).

### Synaptic organization of neuron types

As observed throughout vertebrates (Rall et al., 1966; Pinching and Powell, 1971b, a; Ichikawa, 1976), many neuron types in the zebrafish OB had neurites that were both pre- and postsynaptic to other neurons, with an abundance of reciprocal synapses. This architecture is likely to support divisive normalization (Carandini and Heeger, 2012; Zhu et al., 2013; Roland et al., 2016; Wanner and Friedrich, 2020).

The targeted annotation of intraglomerular synapses between representative neurons revealed subnetworks of GLINs and PLINs that were selectively associated with MCs and RCs. This clear relationship between neuron types, which were classified purely based on anatomical features, and their selective connectivity further validates the taxonomy of cell types. Moreover, the finding that subnetworks associated with MCs and RCs contain distinct, yet structurally similar microcircuits suggests that these subnetworks may perform similar computations in parallel.

Previous results suggested that MCs may inhibit the ruff of RCs via local INs, resulting in anti-correlated odor responses (Zippel et al., 1999), but direct evidence for this hypothesis is lacking. We did not find an obvious synaptic MC◊IN◊RC connectivity motif that may mediate such lateral inhibition although more complex synaptic pathways may exist. Interestingly, dendrites of MCs and RCs formed large physical contacts, which may mediate non-synaptic interactions between neurons. In the zebrafish OB, metabotropic glutamate receptors have been implicated in inhibitory interactions between neighboring PNs (Judkewitz, 2005), raising the possibility that interactions between MCs and RCs may involve paracrine mechanisms. Tight physical associations between neurites were also observed between other neuron types (RC-GLIN1, RC-GLIN4), indicating that non-synaptic interactions may also be involved in additional microcircuits. Moreover, gap junctions were observed between RCs and perinest cells (GLIN1s) and MCs and mixed-synapse cells (GLIN2s) in goldfish (Kosaka and Hama, 1982a), and between MCs and INs in the zebrafish OB (Zhu et al., 2013).

MCs and RCs were both engaged in a connectivity motif that included the principal neurons (MC or RC), a reciprocally connected PLIN (PLIN3v or PLIN1, respectively), and a unidirectionally connected GLIN (GLIN2 or GLIN1, respectively) that was postsynaptic to the PN and projected to extraglomerular targets (Fig. 19). We speculate that this microcircuit normalizes the activity of the principal neuron’s activity via reciprocal inhibition and modifies the activity in other processing channels via unidirectional lateral inhibition or disinhibition. Hence, we expect that the comprehensive structural analysis of neuronal circuits in the OB will promote insights into mechanisms of information processing in olfaction.

## Methods

### Image pre-processing and alignmen

The SBEM volume was acquired previously as described (Wanner et al., 2016). Briefly, images were acquired using a scanning electron microscope (Zeiss Merlin with Gemini II column) equipped with an automated ultramicrotome inside the vacuum chamber (Gatan 3view) (Denk and Horstmann, 2004). Between cuts, partially overlapping rectangular images (“tiles”; n = 154’880) were acquired to generate large composite images of each block face (“sections”; n = 3’918). An initial alignment of tiles and sections was performed using translations based on cross-correlation in the Fourier domain (Preibisch et al., 2009). Visual inspection and computation of deformation fields revealed that this pre-aligned stack occasionally contained non-linear distortions that were typically limited to a few sections and often affected only subsets of tiles. To correct for these distortions, an elastic transformation was applied to sequences of distorted tiles delimited by unaffected, fixed “keyframes” surrounding the distorted tiles. We used the “elastix” framework to correct for these distortions (Klein et al., 2010; Shamonin et al., 2014). After alignment, the final stack was trimmed to a cuboid of 288 x 173 x 98 μm^3^. Contrast was normalized by histogram equalization.

A semantic segmentation model to classify neuropil, somata, sensory axons, blood vessels, extracellular space, and other structures (e.g., surrounding resin) was trained based on manual annotations of image data throughout the stack.

### Neuron reconstruction

Automated image segmentation was performed using flood-filling networks starting from seeds in the neuropil (Januszewski et al., 2018). Networks were trained on expert annotations of three subvolumes (640 x 640 x 100 or 512 x 512 x 128 pixels) in different layers of the OB. A dynamical local realignment procedure was applied during inference (Li et al., 2020) except when large distortions were encountered between sections. Separate network models were trained for the full resolution (9 x 9 x 25 nm) and for a dataset that was downsampled in 2D to 18 x 18 x 25 nm. The base segmentation represents the oversegmentation consensus of segmentations at the full and downsampled resolution in standard and reversed seed order, respectively.

Network models were selected from a pool of versions by comparing the segmentation to skeleton reconstructions of complete neurons distributed throughout the volume. 10 PNs and 11 INs were reconstructed manually by an expert and subsequently refined during comparison to the segmentation. An additional 29 skeletons were generated by external annotators (www.ariadne.ai) using the CORE procedure with two-fold redundancy (Wanner et al., 2016), followed by additional expert curation. A set of 33 skeletons with a total length of 121.5 mm was used for segmentation evaluation.

Volumetric reconstructions of skeletonized neurons (n = 50, including the 33 neurons for model selection) were generated by intersecting skeletons with the agglomeration of the base segmentation and subsequent manual correction of the agglomeration. Additional neurons were reconstructed by direct manual proofreading of the agglomeration. Custom tools (Moenig, 2020a) to correct split and merge errors in the agglomeration were written in Python, accessed the agglomeration via the Brainmaps API (Moenig, 2020b), and used neuroglancer (Maitin-Shepard, 2020) for visualization.

We initially proofread putative PNs (n = 21), starting from somata in the GL. We then identified synaptically connected INs, proofread subsets of the INs with somata contained in the EM volume, and selected synaptic partners of these INs for further proofreading. A second set of neurons (n = 438) was selected pseudo-randomly by reconstructing additional neurons throughout all layers to increase the number of representatives of all neuron classes.

### Coarse reconstruction

The coarse proofreading of PNs involved the removal of all obvious mergers and the fixing of splits to a degree that revealed the overall dendritic morphology of the cell. This procedure included correcting splits in major branches, carefully scanning the outline of the dendrites, and adding all split branches found, but not thoroughly tracing along all the branches. The goal of these procedures was to obtain reconstructions that are sufficiently precise to determine the morphological subtype and to determine the spatial extent of the dendritic tree.

### Morphological feature extraction

PNs were identified based on known morphological features, particularly a myelinated axon, a ruff, and a characteristic tufty dendrite (Kosaka and Hama, 1979, 1982a; Fuller and Byrd, 2005; Fuller et al., 2006a; Braubach and Croll, 2021). Other cells were considered INs. 32 PNs (8 small MCs, 4 intermediate MCs, 12 large MCs, 8 RCs) with CD < 3 were selected for further analysis to extract morphological features, Volumetric reconstructions were transformed into skeletons using the *TEASAR* algorithm (Sato et al., 2000). As this algorithm tends to fill the volume of larger neurites with short appendages, the initial skeletonization was followed by three successive pruning steps to remove short terminal branches of length <100 nm, <400 nm, and <1000 nm, respectively. In addition, a semi-manual procedure was used to generate consolidated representations of somata. This procedure involved the manual definition of a node representing the soma center, the fit of an ellipsoid to the soma volume, the collapse of all segments within the ellipsoid to the center node, and two successive pruning steps to remove remaining short terminal branches outside the ellipsoid (<500 nm and <2500 nm, respectively). Skeletons were represented by directed graphs rooted to the soma center. The total length was calculated as the sum of all branch lengths. The tip ratio was calculated as fraction of the total skeletal length that was contributed by terminal branches. The soma volume was calculated based on the number of voxels belonging to a neuron’s volumetric representation within a sphere adjusted to the soma diameter. This measure may include the volume of branches and spines belonging to a given neuron if these were inside the sphere.

### Neuron classification

Manual neuron classification was performed by five experts (neuroscientists) using a visual user interface based on neuroglancer (Maitin-Shepard, 2020) that allowed for efficient translations, rotations and zoom operations in 3D. Neurons could be viewed individually or in arbitrary combinations to perform comparisons. Users were instructed to group neurons into up to 20 morphological classes without any constraints on features, including the option to leave a cell unclassified if it could not be confidently assigned to a group. The classification of each user was represented as an undirected graph in which each neuron represented a node. If two neurons were assigned to the same class an edge of weight 1 was created. A consensus graph was created by summing up the edge’s weights between all pairs. The result was visualized ordered by the morphological classification presented here.

### Synapse detection

2589 synapses were manually annotated in 128 PN neurites and 232 IN neurites. The cell interior was screened at membrane contact sites between a given pair of cells for the presence of chemical synapse structures, namely a vesicle cloud in the presynapse, vesicles associated with the plasma membrane, and a close contact between the pre- and postsynaptic membrane in at least three consecutive sections. OB neurites are characterized by an extensive presence of vesicles and organelles of the endomembrane system that may be mistaken for synaptic structures. To minimize misclassifications, contact sites were carefully inspected in all three orthogonal viewports in neuroglancer and rated by confidence for the presence of a synapse on a scale of one to five. Here, only synapses with a confidence score of three or higher were included in the analysis.

### PN-IN connectivity within a glomerulus

With the exception of GLIN1s, GLIN2s, GLIN4s and PLINs the connectivity between INs of a given class and PNs was assessed as follows. In pairwise overlays of an IN with all reconstructed PNs of a given glomerulus, each contact site was screened for the presence of synapses. Connections were counted only if the confidence of a synapse was >2. For each IN subtype one to seven examples were inspected (table 1, Fig. 20A). IN – glomerulus pairs were chosen based on the IN passing a dendrite through the glomerular neuropil. Note that this procedure may underestimate connectivity between INs and RCs because the ruff resides outside this volume.

Because GLIN1 dendrites did not enter the glomerular neuropil dendro-dendritic GLIN1-mitral cell connections were improbable. The axo-dendritic connectivity was assessed by pairwise screening for connections between GLIN1s and PNs of the glomerulus innervated by the GLIN1 axon (629 pairs). For GLIN4, pairwise overlays with PNs from both glomeruli innervated by the GLIN4 dendrite and axon were screened (330 GLIN4-MC pairs, 56 GLIN4-RC pairs). For GLIN2 and PLIN1, IN-MC connectivity was assessed as described above. Because dendrites of GLIN2s and RCs are a purely postsynaptic, axo-dendritic or axo-axonic connectivity was specifically analyzed throughout the dataset in pairwise overlays of 19 GLIN2 and 17 RCs. To assess the connectivity between RCs and PLIN1s, pairwise overlays between all PLIN1s and all RCs whose ruff neuropil was innervated by a PLIN1 dendrite were screened for synapses. The number of cells and the number of glomeruli analyzed for each IN subclass are summarized in table 2.

### Sensory input

Sensory axons were excluded from the instance segmentation of neurons in our dataset. To search for synaptic contacts between INs and sensory axons the 3-dimensional visualization of a semantic segmentation of all sensory axons was overlayed with one IN at a time and the dendritic branches of the IN were followed to identify contacts with the sensory axons.

## Acknowledgements

We thank Jeremy Maitin-Shepherd, Markus Rempfler, and Tim Blakely for their generous technical advice and constructive feedback which was crucial for the successful implementation of the proofreading tool. We are grateful to the Friedrich group for insightful discussions. This work was supported the by Novartis Research Foundation, by the European Research Council (ERC) under the European Union’s Horizon 2020 research and innovation program (grant agreement no. 742576), by the Human Frontiers Science Program (grant no. RGP0015/2010), and by the Swiss National Science Foundation (grants no. CRSII3_130470/1, 31003A_172925/1, 310030_219500, 31003A_152833/1, 310030_212236).

## Author Contribution

**N.R.M.**: Conceptualization, Methodology, Investigation, Data curation, Visualization, Validation, Software, Formal analysis, Writing—original draft, Writing—review & editing, Created training data for segmentation and tissue classification models, Developed software for neuron reconstruction, classification and data analysis, Reconstructed all neurons, Deep classification of neurons, Synapse annotation. Contributed equally with M.J.. No competing interests declared.

**M.J.**: Software, Validation, Methodology, Data curation Developed and applied segmentation and tissue classification. Contributed equally with N.R.M.. M.J. is an employee of Google.

**S.G.**: Software, Data curation, Validation, Writing—review & editing, Aligned the EM dataset. No competing interests declared.

**B.H.**: Investigation, Writing—review & editing, Neuron classification. No competing interests declared

**N.Z.T**: Investigation, Neuron classification. No competing interests declared**R.E.M.C.**: Investigation, Neuron classification. No competing interests declared.

**T.M.**: Investigation, Writing—review & editing, contributed to EM data acquisition. No competing interests declared.

**A.A.W.**: Investigation, Writing—review & editing, contributed to EM data acquisition; No competing interests declared.

**C.G.**: Investigation, contributed to EM data acquisition; No competing interests declared

**R.W.F**: Conceptualization, Supervision, Funding acquisition, Writing—original draft, Writing— review & editing, Investigation, Neuron classification. No competing interests declared.

